# Flaviviruses Hijack HuR to Suppress CNPY3: A Novel Pathway for Immune Evasion and Therapeutic Intervention

**DOI:** 10.1101/2024.12.31.630865

**Authors:** Xiaoyan Ding, Jiuxiang He, Jing Zhao, Yuxin Zhou, Xiaozhong Chen, Jun Tong, Wenxuan Quan, Minyue Qiu, Dong Hua, Minchi Liu, Shan Guan, Jintao Li

## Abstract

Flaviviruses represent a serious global health threat, infecting millions annually with high rates of cross-infection among different flaviviruses, yet no broad-spectrum antiviral therapy currently targets these pathogens. Through biological big data analysis, we observed a significant downregulation of Canopy FGF Signaling Regulator 3 (CNPY3), a positive regulator of defense responses, in flavivirus-infected patients, with the level of downregulation correlating with infection severity. This observation was validated in cellular and animal models. Mechanistically, we demonstrate that flaviviruses hijack the RNA-binding protein Human antigen R (HuR) to destabilize CNPY3 mRNA via adenine-uridine-rich elements (AREs), thus downregulating CNPY3 expression and promoting viral replication. Our findings identify CNPY3 as a novel antiviral mediator essential for the TLR-mediated type I interferon (IFN-I) pathway in defense against flaviviruses. In vitro, CNPY3 overexpression reduced DENV and ZIKV replication, confirming its antiviral role. To explore therapeutic potential, we developed lipid nanoparticles (LNPs) encapsulating CNPY3 mRNA (LNP-CNPY3), and early administration in flavivirus-infected mice improved survival, reduced viral loads, and attenuated neuroinflammation by enhancing antiviral and interferon responses. Together, these findings establish CNPY3 as a critical antiviral target and support LNP-CNPY3 as a promising therapeutic approach against flavivirus infections.

## Introduction

Flaviviridae are a significant group of human pathogens, including dengue (DENV), West Nile (WNV), Zika (ZIKV), and yellow fever viruses (YFV). These viruses are primarily transmitted through mosquito bites, exhibit high pathogenicity, and have a broad host range, making them prone to large-scale outbreaks^1,2^. Currently, therapeutic options for flavivirus infections are limited, primarily relying on supportive care and vaccine prophylaxis. However, the efficacy of vaccines tends to be virus-specific, which emphasizes the need for broad-spectrum antiviral drugs, especially for patients infected with flaviviruses or at risk of repeated exposure to flaviviruses^3^.

To develop effective therapies against flaviviruses, it is crucial to gain a comprehensive understanding of virus-host interactions and the mechanisms underlying the antiviral immune response ^4^. The innate immune system serves as the first line of defense against viral infections and plays a critical role in initiating and shaping the adaptive immune response. Pattern Recognition Receptors (PRRs), such as Toll-like receptors (TLRs), are essential components of the innate immune system that recognize Pathogen-Associated Molecular Patterns (PAMPs) and activate signaling pathways leading to the production of antiviral proteins, including type I and type III interferons (IFNs))^5–7^. TLRs, in particular, detect a variety of PAMPs and signal through Mitochondrial Antiviral Signaling Protein (MAVS) to activate transcription factors such as IRF3 and NF-κB, which are key regulators of type I IFN production^8–10^. Conversely, flaviviruses have evolved sophisticated strategies to evade host immune detection and inhibit antiviral responses. They achieve this by concealing their molecular signatures and modulating the TLR signaling pathway at multiple levels, ultimately promoting viral replication and survival within the host^11–14^. Despite these insights, there remains a significant gap in identifying optimal molecular targets for antiviral therapies, particularly those that are broad-spectrum and effective against multiple flaviviruses. One promising approach to identifying new antiviral targets involves the screening and analysis of large-scale transcriptomic data from clinical patient samples, which could reveal novel molecules involved in the host’s antiviral defense mechanisms.

In this study, we identified CNPY3 as a novel antiviral target that modulates the TLR-mediated type I interferon (IFN-I) signaling pathway, influencing flavivirus replication. Disruption of CNPY3 impairs IFN-I signaling, enhancing viral replication and immune evasion. Both positive and negative-strand RNAs of flaviviruses, the latter being an intermediate in replication, play critical roles in this immune escape mechanism, highlighting their impact on host immune response manipulation. We propose a lipid nanoparticle (LNP)-based therapeutic approach using in vitro transcribed mRNA encoding CNPY3 (Lipid-CNPY3 mRNA) to target flavivirus infections, showing promise as an effective antiviral treatment strategy.

## Results

### CNPY3 Downregulation is Linked to Flavivirus-Infection Severity

Deciphering the host factors essential for the replication of Flaviviruses holds the key to devising efficacious therapeutic interventions. According to analysis DENV-infected dendritic cells (DCs) and DENV infected patients, we found that the type I interferon signaling pathway genes such as *USP18*, *MIX1*, *OAS1*, *OAS2*, *ISG20*, etc., were significantly up-regulated while the positive regulation of defense response, including *FCER1A*, *IFNGR1*, *CD300LF*, *CLEC7A*, *CLEC4A*, *CNPY3*, *TLR5*, and *TGM2* were significantly down-regulated (Fig. 1A,B and Fig. S1). Further analysis in a human monocyte cell line (THP-1) infected with DENV-1, DENV-2, DENV-3, and DENV-4 showed that genes *OAS1, IFI27* and *MX1*were significantly increased, and *CNPY3*, *FCER1A* and *TLR5* were significantly decreased after infection with any of the four DENV serotypes (Fig.1C). Interestingly, the expression of immune response genes *CNPY3*, *FCER1A* and *TLR5* decreased, especially *CNPY3* which was never reported before upon flavivirus infection. To explore if the expression of *CNPY3* could also be inhibited by other flaviviruses, the expression of *CNPY3* after ZIKV infection was measured. ZIKV is another member of the clinically important family of mosquito-borne Flaviviruses ^15,16^. Consistent to our suspicion, *CNPY3* expression was also significantly inhibited during ZIKV infection (Fig.1D and E).

**Fig. 1.**
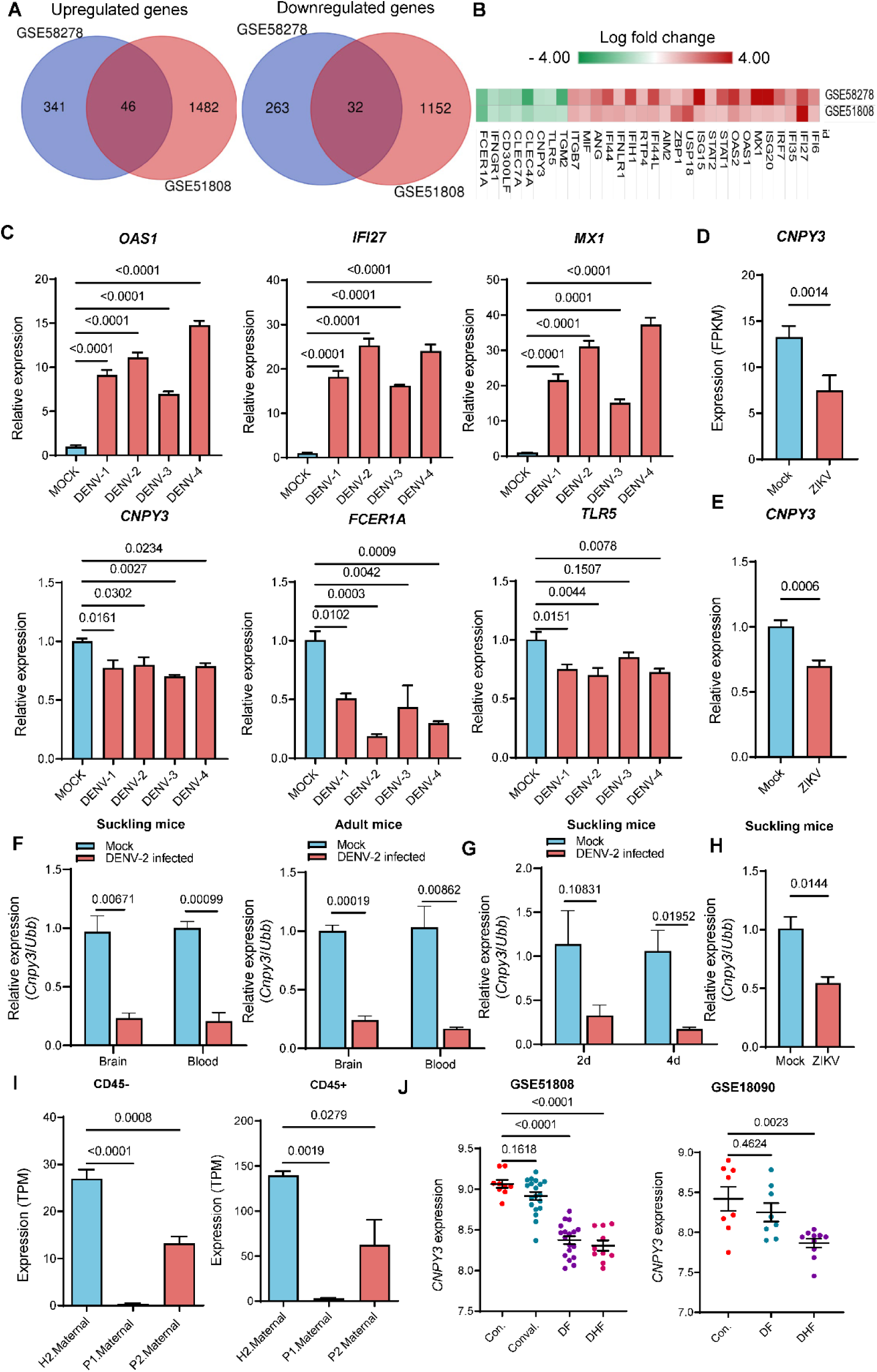
*CNPY3* expression is repressed and negatively correlated with disease severity during flavivirus infection. (A and B) Transcriptome analysis of DENVs infected human monocyte-derived dendritic cells (GSE58278) and DENV patients (GSE51808). Venn diagram representing data sets sharing the number of common DEGs (A). Heat map showing consistently expressed genes in GSE58278 and GSE51808 (B). (C) Expression changes of key genes determined by qPCR in DENV-infected THP-1 cells (MOI = 2). Gene expression in mock-infected cells as the control. Adjusted P values were determined by a one-way ANOVA followed by Dunnett’s multiple comparisons test. (D and E) *CNPY3* expression re-analysis of RNA-seq data (GEO accession GSE80434) in human neural progenitor cells (hNPCs) after African ZIKV infection (D) and determined by qPCR in ZIKV-infected A549 cells (MOI = 2, E). *CNPY3* expression in mock-infected cells as the control. P values were determined by an unpaired two-tailed t-test. (F and G) Analysis of *Cnpy3* expression levels in brain and blood of suckling mice and A6 adult mice (F) and *Cnpy3* expression in suckling mice after DENV-2 intracranial infected 2d and 4d (G). The brain and whole blood of 3-days-old BALB/c mice, and 4-week-old A6 mice were collected for expression analysis after DENV-2 intranasal infected 7 d or 9 d, respectively. To test the expression at different points in time, the whole blood of 3-days-old BALB/c mice after DENV-2 intracranial infected 2 d and 4 d were collected. Mock: controls without DENV-2 virus infection; Infected: DENV-2 virus infection. Results are presented as mean ±S.E.M. (n = 3). P values were determined by a multiple unpaired two-tailed t-test. (H) *Cnpy3* expression in suckling mice after ZIKV intracranial infected 4d. Mock: controls without ZIKV infection; ZIKV: ZIKV infection. Results are presented as mean ±S.E.M. (n = 3). P values were determined by an unpaired two-tailed t-test. (I) *CNPY3* expression in RNA-seq data (GEO accession GSE139181) of placental tissue from ZIKV-infected pregnant women compared to healthy controls (n=3 tissues per patient). Placenta disc was divided into CD45+ immune and CD45- non-immune fractions. H2. Maternal, tissues from the healthy placental; P1. Maternal and P2. Maternal tissues from two individual patients who were infected with ZIKV. Adjusted P values were determined by a one-way ANOVA followed by Dunnett’s multiple comparisons test. (J) *CNPY3* expression in DENV patients with different disease progression. Raw data came from GSE18090 and GSE 51808. Each dot presented *CNPY3* expression values from one patient. con., healthy person; conval., convalescent patient; DF, dengue fever; DHF, dengue hemorrhagic fever. Adjusted P values were determined by a one-way ANOVA followed by Dunnett’s multiple comparisons test.

To validate our findings, we further analyzed *Cnpy3* expression in *in vivo* Flavivirus-infected mice models. Consistent with the *in vitro* findings, we found *Cnpy3* expression was also significantly down-regulated in brain and blood after DENV-2 infection in both suckling and adult mouse models (Fig. 1F). In addition, the degree of inhibition of *Cnpy3* expression in the blood of mice was higher as the duration of infection was prolonged (Fig.1G). Similarly, *Cnpy3* was also downregulated in the blood of mice after ZIKV infection (Fig. 1H). Furthermore, clinical data indicate that the expression of *CNPY3* is significantly suppressed in patients infected with DENVs or ZIKV, with greater suppression observed in cases of severe infection cases (Fig. 1I and J). These results imply that *CNPY3* expression was repressed during Flavivirus infection and negatively correlated with the severity of disease.

### Flaviviruses Hijack HuR to Suppress CNPY3 Expression

Previous studies have shown that CNPY3 is post-transcriptionally regulated by the RNA-binding protein HuR, which stabilizes CNPY3 mRNA by directly binding to it ^17^. Consistent with this, our experiments confirmed that HuR interacts with both the ORF and 3’ UTR regions of CNPY3 mRNA (Fig. S2A and S2B). Silencing HuR in U937 monocytic cells led to a significant decrease in CNPY3 expression (Fig. S2C), suggesting that disruption of HuR binding impairs CNPY3 stability. We hypothesize that flavivirus RNAs hijack HuR, which normally stabilizes CNPY3 mRNA, leading to decreased CNPY3 expression. To validate this hypothesis, we analyzed potential interactions between HuR and the RNA of several flaviviruses, including Yellow Fever, Dengue, Zika, West Nile, and Japanese Encephalitis viruses, using RPISeq. The analysis indicated a high likelihood of HuR interacting with both the positive and negative RNA strands of these viruses (Table S1).

Then we examined the localization of viral RNA, CNPY3 mRNA, and HuR in normal and DENV-2-infected RAW264.7 cells. In uninfected cells, CNPY3 mRNA was found in both the nucleus and cytoplasm, while HuR primarily localized to the nucleus, co-localizing with CNPY3 mRNA (Fig. 2A). After DENV-2 infection, viral RNA accumulated around the nuclear periphery, and HuR translocated to this region, co-localizing with DENV-2 RNA, while CNPY3 mRNA localization remained unchanged (Fig. 2B). This HuR translocation was even more pronounced in brain cells from infected mice (Fig. 2C and D). HuR is known to bind adenine-uridine-rich elements (AREs)^18,19^. We further analyzed the positive and negative RNA strands of various flaviviruses and found multiple AREs in both strands (Fig. S3). RNA pull-down experiments demonstrated a strong interaction between HuR and the 5’ UTR of the DENV-2 negative-strand RNA, which contains several AREs, while the control positive-strand RNA without AREs showed no significant binding (Fig. 2E). RNA immunoprecipitation (RIP) further confirmed HuR binding to the 5’ UTR of the DENV-2 negative-strand RNA (Fig. 2F).

**Fig. 2.**
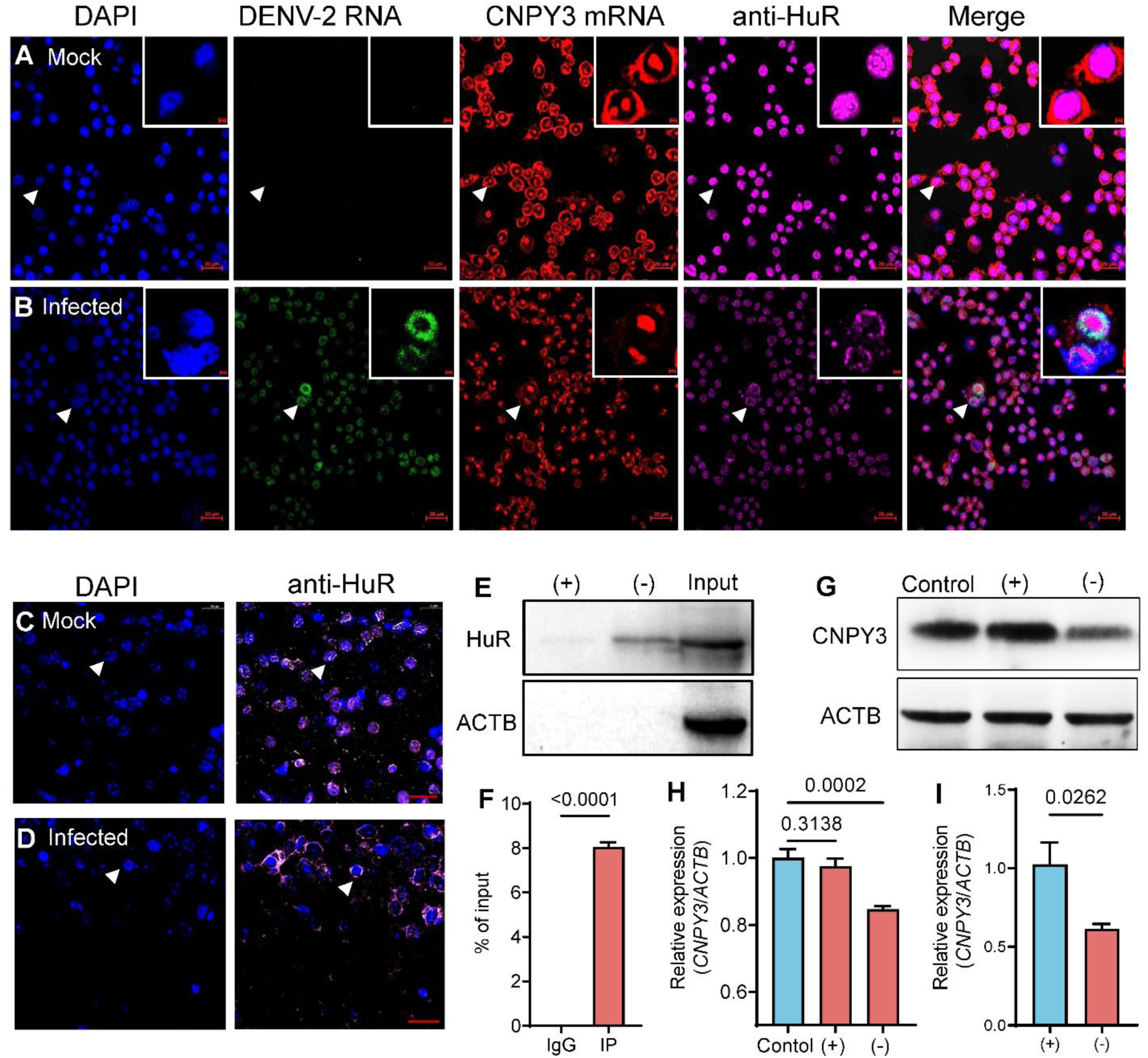
DENV-2 RNA hijacks HuR and inhibits CNPY3 expression. (A and B) Location analysis of DENV-2 RNA, CNPY3 mRNA, and HuR protein in Mock-infected (A) and DENV-2-infected (B) monocyte/macrophage-like cell line RAW264.7. At 48 hpi, RAW264.7 cells were FISH analysis using a DENV-2 RNA probe (green) or a CNPY3 mRNA probe (red) and stained with DAPI (blue), anti-HuR antibody (pink). Bar = 20μm. (C and D) HuR protein location in mock-infected (C), infected with DENV-2 (D) in mice brain. 3d-old-suckling mice brains infected with DENV-2 for 6 d, brain tissues were fixed and stained with DAPI- and HuR-specific antibodies and analyzed by Pannoramic DESK digital slice scanner. Bar = 20μm. (E) RNA-pulldown was used to detect the interaction between DENV-2 minus strand 5’UTR and RNA binding protein HuR. (F) RIP analysis was used to measure the association of HuR with DENV-2 minus strand 5’UTR by using either anti-HuR antibody or control IgG. Results are presented as mean ±SEM (n = 3). P values were determined by an unpaired two-tailed *t*-test. (G and H) Protein and mRNA expression level of CNPY3 in U937 cells treated with DENV-2 minus strand 5’UTR and DENV-2 3′UTR. Results are presented as mean ±SEM (n = 3). Adjusted P values were determined by a one-way ANOVA followed by Dunnett’s multiple comparisons test. (I) DENV minus strand represses the expression of CNPY3 in overexpression HuR 293T cells. Results are presented as mean ±SEM (n = 4). P values were determined by an unpaired two-tailed *t*-test. (+), 3′UTR of DENV positive strand; (−), 5′UTR of DENV minus strand; Input, positive control; Control, *in vitro* transcription (IVT)- eGFP mRNA.

Transfecting cells with the 5’ UTR of the DENV-2 negative-strand RNA led to a significant reduction in CNPY3 expression, while the control RNA without HuR binding sites did not alter CNPY3 levels (Fig. 2G and H). Moreover, in HuR-overexpressing cells, transfection with DENV-2 negative-strand RNA caused a notable decrease in CNPY3 expression compared to control RNA transfection (Fig. 2I). These findings confirm that flavivirus RNA hijacks HuR via AREs, thereby repressing CNPY3 expression and contributing to immune evasion.

### CNPY3 modulates robust antiviral responses via TLR-mediated innate immunity

Toll-like receptors (TLRs) play a crucial role in host defense and innate immune responses against microbial invasions, triggering signaling events that lead to the production of type I interferons and the establishment of antiviral immunity^20^. CNPY3 is essential for the proper folding and expression of most TLRs, except TLR3^21^. Consistent with previous findings, we observed a positive correlation between CNPY3 expression and the expression of several TLRs—including TLR1, TLR2, TLR4, TLR5, TLR6, TLR7, TLR8, TLR9, and TLR10 in the whole blood of healthy individuals (Fig. S4). This positive correlation was also evident in patients infected with DENV (Fig. S5). The cell and tissue tropism of flaviviruses significantly influences the outcomes of infection^22^, with monocytes and dendritic cells (DCs) being primary targets for flavivirus entry ^23,24^. Notably, CNPY3 is highly expressed in the brain, blood, bone marrow, and lymphoid tissues (Fig. S6A). Single-cell sequencing data further revealed that CNPY3 is most abundantly expressed in dendritic cells, endothelial cells, and monocytes within the blood—cells that are key targets of flavivirus infection (Fig. S6B and S6C). Given its high expression in these critical immune cells and its role in facilitating the proper folding and function of TLRs, CNPY3 likely serves as an important barrier against flavivirus infection.

To confirm the antiviral role of CNPY3, we first constructed a CNPY3 overexpression vector and transfected it into cells with low endogenous CNPY3 expression, effectively increasing CNPY3 levels (Fig.3 A and B). We then evaluated the susceptibility of these CNPY3- overexpressing cells to flavivirus infections, including DENV and ZIKV. The results showed a significant reduction in DENV-2 and ZIKV replication in CNPY3-overexpressing 293T cells (Fig. 3C, D, E, F, and G). Similarly, ZIKV infection was also markedly reduced in CNPY3- overexpressing A549 human lung adenocarcinoma cells compared to cells transfected with an empty vector (Fig. 3H, I). These findings demonstrate that CNPY3 plays a key role in the antiviral response against flavivirus infections.

**Fig. 3.**
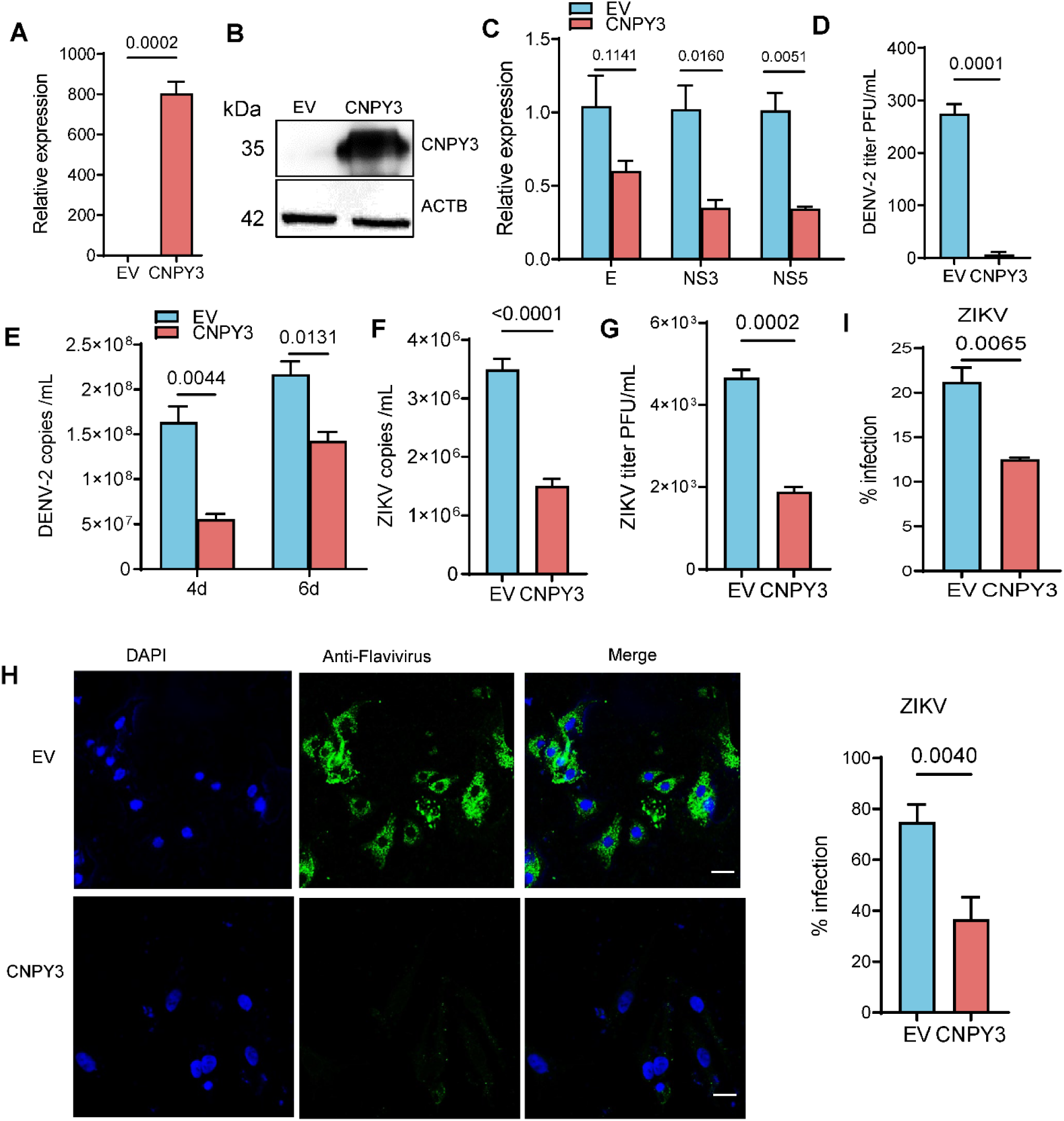
CNPY3 overexpression represses flavivirus replication *in vitro*. (A and B) mRNA and protein expression level of CNPY3. HEK 293T cells were transfected with pCDNA3.1 (EV) or pCDNA3.1-CNPY3 (CNPY3). After 36 h, CNPY3 expression was analyzed by (A) RT-PCR and (B) western blot. Results are presented as mean ±SEM (n = 3). P values were determined by an unpaired two-tailed t-test. (C and D) Expression levels of DENV-2 E, NS3 and NS5 gene (C) in HEK 293T cells and (D) quantification by plaque assay of infectious virus particles in the supernatants. HEK 293T cells were transfected with equal concentrations of both the plasmids. After 36 h, Cells were infected with DENV-2 at an MOI of two. 18 h later, cells were collected for mRNA relative expression and the cellular supernatant was collected for plaque assay. Results are presented as mean ±SEM (n = 3). P values were determined by multiple t-test. (E) DENV-2 copy number in HEK 293T cell supernatant at different time point. HEK 293T cells were transfected with equal concentrations of both the plasmids. Cellular supernatant was collected at 4d and 6d for virus RNA extraction and qPCR assay. Results are presented as mean ±SEM (n = 3). P values were determined by multiple t-test. (F and G) ZIKV copy number and virus titers in HEK 293T cell supernatant. HEK 293T cells were transfected with equal concentrations of both the plasmids. After 36 h, Cells were infected with ZIKV at an MOI of two. 24 h later, the cellular supernatant was collected for virus RNA extraction and qPCR assay (n = 6) and plaque assay (n = 3). Results are presented as mean ±SEM. P values were determined by an unpaired two-tailed t-test. (H) Immunofluorescence observation and quantified of ZIKV infection with A549 cells. Bar = 20 μm. Results are presented as mean ±SEM (n = 8). (I) ZIKV infection analysis by flow cytometry. Results are presented as mean ±SEM (n = 3). A549 cells were transfected with equal concentrations of EV or CNPY3. 24h later, cells were infected with ZIKV at an MOI of two. The cells were fixed and stained by flavivirus group antibody at 48hpi. Three immunofluorescent slices per group have been observed. Quantitative analyses were calculated for a total of eight regions of the three slices. The cells were also collected and stained for flow cytometry. P values were determined by an unpaired two-tailed *t*-test.

Based on our findings, we hypothesized that flaviviruses evade host immune surveillance by inhibiting CNPY3 expression. To test this, we suppressed CNPY3 expression in THP-1 cells (Fig. 4A and 4B) and examined the expression of genes involved in the type I interferon (IFN-I) antiviral signaling pathway, which is the first line of defense against flaviviruses^25^. As expected, the production of type I interferon, specifically IFN-β, was reduced in CNPY3-downregulated THP-1 cells (Fig. 4C). Similarly, interferon-stimulated genes (ISGs) such as *MX1*, *OAS1*, *ISG15, OAS2*, and *USP18* were also downregulated (Fig. 4D). Next, we analyzed DENV-2 infection in CNPY3-knockdown THP-1 cells. The results showed higher expression of DENV-2 genes *E*, *NS3*, and *NS5* in CNPY3-knockdown cells compared to controls (Fig. 4E). Western blot analysis also revealed increased accumulation of the E protein in CNPY3-knockdown cells (Fig. 4F). Moreover, these cells exhibited a higher viral load, particularly at the later stages of infection (6 days post-infection), with a significantly greater accumulation of DENV-2 copies compared to control cells (Fig. 4G). These results confirmed our hypothesis that flaviviruses may disrupt the CNPY3-regulated TLR-mediated IFN-I antiviral signaling pathway by downregulating CNPY3 expression. This inhibition enables successful infection of host cells and allows flaviviruses to evade host immune surveillance.

**Fig. 4.**
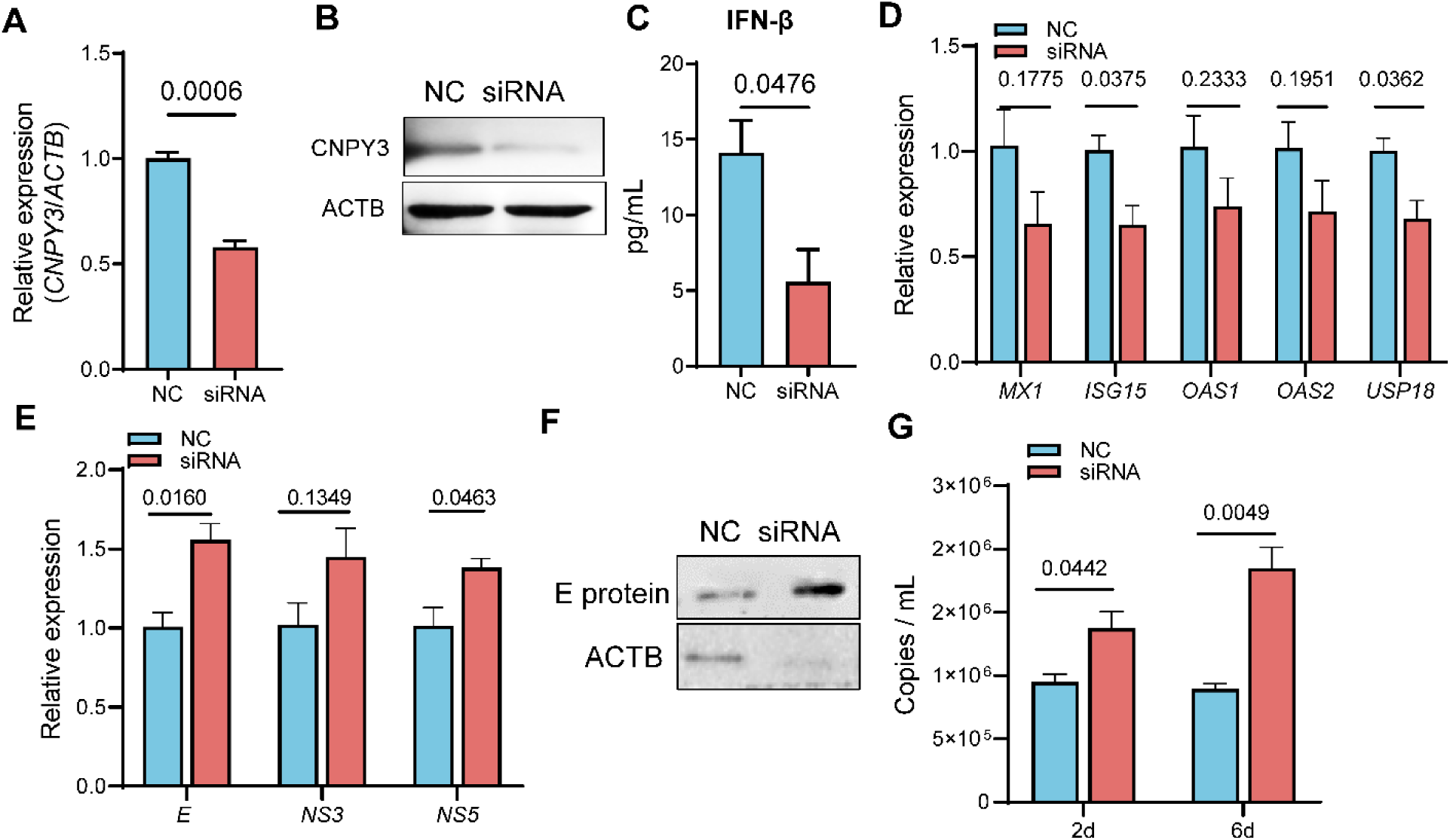
Inhibition of CNPY3 expression disrupts the IFN-I antiviral signaling pathway and promotes viral infection. (A) mRNA and (B) protein expression level of CNPY3 in transfected THP-1 cells. (C and D) Determine IFN-β content (C) and analysis ISGs genes expression in THP-1 cells. THP-1 cells were transfected with a CNPY3-siRNA or Negative Control (NC)-siRNA. After 36 h, RNA and protein were then extracted for analysis. (E and F) E, NS3 and NS5 RNA expression level (E) and E protein expression level (F) in THP-1 cells after DENV-2 infected. THP-1 cells were transfected with CNPY3-siRNA or negative control (NC)- siRNA. 36 h later, transfected cells were infected with DENV-2 at multiplicity of infection (MOI) of 2 for 18 h. The cells RNA and protein were then extracted and analyzed. (G) DENV-2 copies in THP-1 cells supernatant at different times. NC, THP-1 cells were transfected Negative Control (NC) siRNA. siRNA, THP- 1 cells were transfected with a CNPY3 siRNA. *ACTB* was an internal reference gene. Results are presented as mean ±S.E.M. (n = 3). P values were determined by an unpaired two-tailed *t*-test.

### Evaluation of LNP-CNPY3 as a Therapeutic Agent against Flavivirus Infections

To assess the potential of CNPY3 as a therapeutic agent against flaviviruses infection, we engineered nanoparticles containing *Cnpy3* mRNA with modified 5’UTR, 3’UTR, and poly(A) tail. The average particle sizes of LNP and LNP-*Cnpy3* were approximately 100 nm (Fig. S7A). TEM analysis revealed well-defined spherical particles with a zeta potential ranging from −1mV to −3 mV (Fig. S7B and Fig. S7C), indicating favorable characteristics for delivering CNPY3 in therapeutic applications.

We next explored the effectiveness of LNP-*Cnpy3* in a mouse model. LNP-*Cnpy3* or empty LNP was intracarotid administered at various stages—before, during, and after lethal DENV-2 infection—and survival rates were assessed. Administering LNP-*Cnpy3* at the early stage of DENV-2 infection significantly increased mouse survival, while late-stage administration had little effect on survival (Fig. S8). To investigate the molecular mechanisms, we conducted transcriptome analysis on brain samples from suckling mice treated with LNP-*Cnpy3* or LNP at the early stage (1day post-infection, dpi). LNP-*Cnpy3* treatment led to the upregulation of 149 genes and downregulation of 64 genes compared to LNP treatment (Fig. S9A). Many of the upregulated genes were associated with the interferon-beta response (Fig. S9B), indicating activation of the IFN-beta signaling pathway. Ingenuity Pathway Analysis (IPA) further revealed that genes affected by LNP-*Cnpy3* treatment have the potential to suppress flavivirus replication (Fig. 5A) and activate several antiviral pathways, such as hyperchemokinemia in influenza pathogenesis, interferon signaling, and activation of IRF by cytosolic pattern recognition receptors (Fig. 5B). Cnpy3 upregulation was linked to increased expression of transcriptional programs indicative of IFN-I signaling, including activation of *IFNB1*, *STAT1*, *IRF3, IRF7, IFNG, NFKB1* and *TLR2* (Fig. 5C, D and Fig. S10). These findings were corroborated by ELISA and RT-PCR analyses of brain samples, which showed that LNP-*Cnpy3* treatment promoted the production of Ifn-α and Ifn-β (Fig. 5E, F). Additionally, the expression of interferon regulatory factor 7 (*Irf7*) and several ISGs, including *Mx1*, *Isg15*, *Oas1*, *Oas2*, and *Usp18*, sharply increased after LNP-*Cnpy3* treatment (Fig. 5G-J). These results suggest that LNP-*Cnpy3* promotes the early production of IFN-I and is therefore more effective when administered during the early stages of flavivirus infection.

**Fig. 5.**
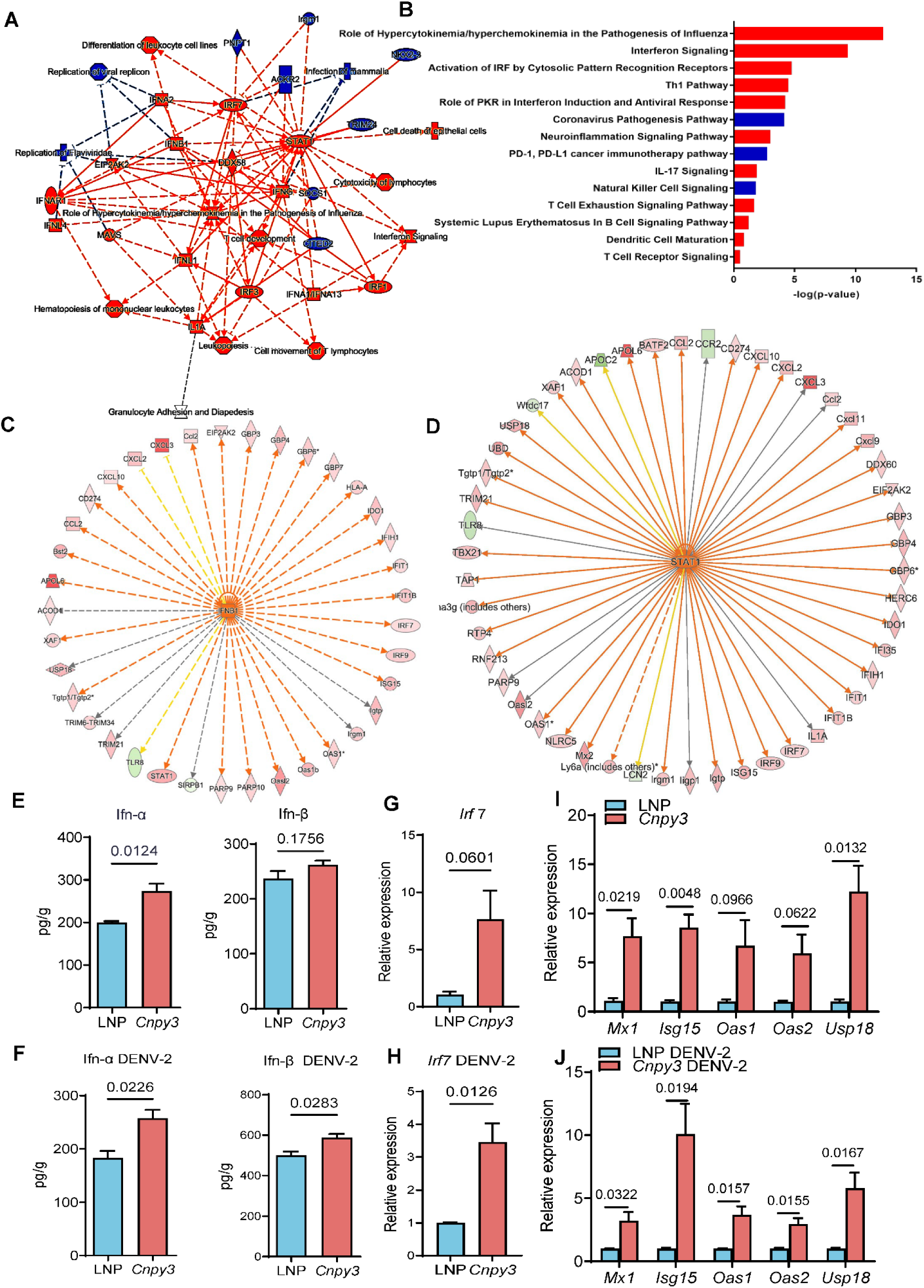
LNP-*Cnpy3* enhanced type I interferon signaling in early-stage of flavivirus infection. (A) Graphical summary of Ingenuity Pathway Analysis (IPA). The blue lines mean inhibited and red lines mean activated. (B) Canonical pathways. The color of the bars indicates whether the pathway is expected to be activated (red bars) or inhibited (blue bars). The length of the bars indicates the significance of overlap between molecules in the dataset and pathways in the QIAGEN knowledge base. Significance calculated based on the right-tailed Fisher exact test and −log(p-value) is shown on the x-axis of the bars. (C and D) IPA identified IFNB1 and STAT1 as upstream regulators. Orange for up-regulation, green for suppression. (E and F) Ifn-α and Ifn-β contents. (G-J) IRF7 and ISGs expressions. 3-day-old mice were infected or uninfected with DENV-2 (60 PFU) and treated with 1μg LNP-*Cnpy3* or LNP at the meanwhile. After 1dpi, brains RNA and proteins were extracted and analyzed. P values were determined by an unpaired two-tailed *t*-test.

We further confirmed the effect of LNP- *Cnpy3* treatment in the early stages of flavivirus infection. Suckling mice were infected with DENV-2 (60 PFU) and treated with 1 μg of LNP- *Cnpy3* or empty LNP simultaneously. LNP- *Cnpy3* treatment significantly reduced DENV-2 NS1 expression (Fig. 6A) and viral RNA copies (Fig. 6B) in the brain at 6 days post-infection (dpi). Encephalitis and encephalopathy are common neurological outcomes following flavivirus infection^26^, with neuroinflammation being initiated by activated microglia. Iba-1 expression, a marker for activated microglia, was notably reduced in LNP- *Cnpy3*-treated mice, indicating decreased neuroinflammation (Fig. 6A). Histopathological analyses at 6 dpi showed that, while LNP-treated mice exhibited damage in the cerebral cortex, the brains of LNP- *Cnpy3*-treated mice maintained intact pia mater, molecular and granular layers, as well as a well-preserved pyramidal neuron layer. Similar results were observed in the hippocampus (Fig. 6C). Transcriptome analysis between LNP and LNP-*Cnpy3*-treated brains revealed 190 upregulated genes and 640 downregulated genes (Fig. S11A). Many of the downregulated genes were associated with inflammatory responses (Fig. S11B). Specifically, genes related to pro-inflammatory responses, such as *Txnip*, *Nlrp3*, *C3ar1*, *Mefv*, *C3*, and interleukin-6 family signaling genes like *Lif*, *Il31ra*, *Il6ra*, *Cntf*, and *Clcf1*, were downregulated in LNP-CNPY3- treated brains (Fig. S11C), indicating reduced production of pro-inflammatory factors and confirming the therapeutic effect of LNP-*Cnpy3* on encephalitis. Consistent with previous findings, LNP-*Cnpy3* treatment delayed the onset of symptoms in DENV-2-infected mice and improved survival rates (Fig. 6D). We further evaluated the efficacy of LNP-*Cnpy3* in other DENV serotypes. Suckling mice infected with DENV-1, DENV-3, or DENV-4 and treated with 1 μg of LNP-CNPY3 or LNP also exhibited delayed onset of infection and higher survival rates. Notably, LNP-CNPY3 treatment against DENV-1 achieved a 100% survival rate (Fig. 6E). To assess the therapeutic potential of LNP-*Cnpy3* against other flaviviruses, one-day-old mice were infected with ZIKV (1000 PFU) and treated with 1 μg of LNP-*Cnpy3* or LNP. LNP- *Cnpy3* treatment inhibited ZIKV replication (Fig. 6F and 6G) and preserved more NeuN-positive cells, indicating that it alleviated ZIKV-induced neuronal damage (Fig. 6F). Additionally, LNP- CNPY3-treated mice displayed reduced brain pathology, evidenced by thicker cortical plates, smaller ventricles, and a higher density of Nissl bodies in their brains, as shown by Nissl staining (Fig. 6H). Collectively, these findings support the role of CNPY3 in inhibiting flavivirus replication and pathogenesis in vivo and highlight LNP-CNPY3 as a promising candidate for therapeutic intervention against flavivirus infections.

**Fig. 6.**
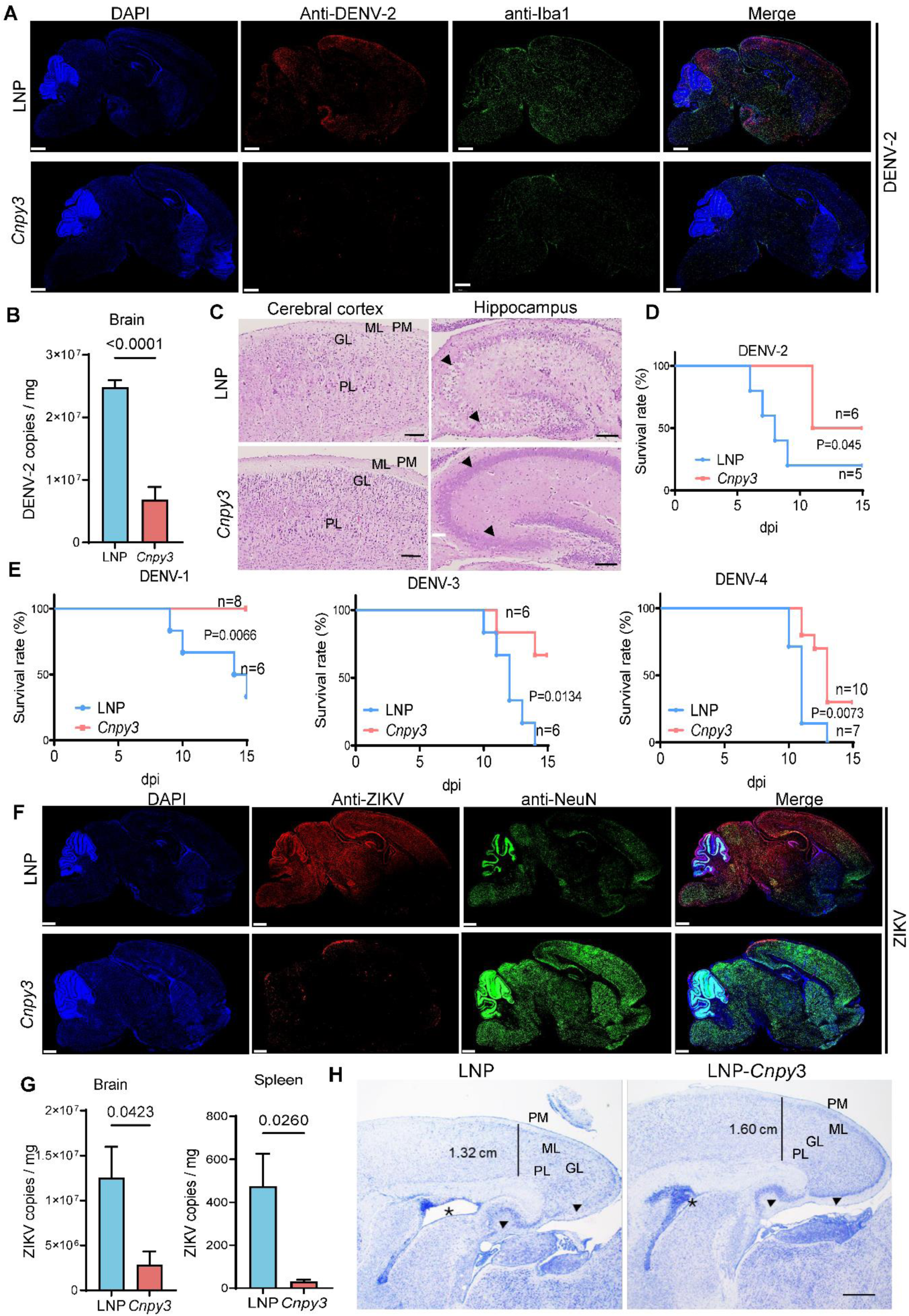
LNP-delivered Cnpy3 mRNA represses flavivirus replication and encephalitis *in vivo*. (A) Immunofluorescence observation. Staining against DENV-2 NS1 or microglia cells was performed using specific antibodies. Nuclei were counterstained with DAPI and imaging was performed using digital slice scanning system. Scale bar = 500 μm. Two immunofluorescent slices per group have been observed. (B) DENV-2 viral load in suckling mice brain. Data represent the mean ±SEM (n = 8), P values were determined by an unpaired two-tailed t-test. (C) Histological analysis of brain. Scale bar = 100 μm. Image is representative of two mice that showed similar results. PM, pia mater; ML, molecular layer; GL, granular layer; PL, pyramidal layer. Damaged neurons (arrows) are frequent in LNP treatment group as compared to LNP-Cnpy3 treatment. 3-day-old BALB/c mice brains were infected with DENV-2 and treated with either LNP-Cnpy3 or LNP alone. On 6 days post-infection (dpi), brains of mice were analyzed. (D and E) Survival rates of the DENV-2 (D) and other DENVs (E) infected mice. 3-day-old BALB/c mice were treated with LNP-*Cnpy3* or LNP by intracranial injection and infected with different DENVs at meanwhile. Survival rates were statistically analyzed using the log-rank (Mantel–Cox) test. (F) Immunofluorescence analysis at 8 dpi of 1-day-old C57BL/6 mice brains infected with ZIKV and treated with either LNP-*Cnpy3* or LNP, staining against flavivirus specific antibody or neuronal nuclei (NeuN) antibody. Nuclei were counterstained with DAPI and imaging was performed using digital slice scanning system. Scale bar = 500 μm. (G) ZIKV copies numbers in suckling mice brain and spleen. Data represent the mean ±SEM (n = 4), P values were determined by an unpaired two-tailed t-test. (H) Nissl histochemical staining observation. Arrows indicated the Nissl bodies. vertical lines indicate cortical thickness and asterisks indicate ventricle dilatation. Scale bar = 500 μm. Image shown is representative of 3 animals that showed similar results.

## Discussion

The innate immune system as the host’s first line of defense against viral infections, with Toll-like receptors (TLRs) playing a critical role in initiating immune responses. CNPY3 is a TLR-specific co-chaperone for HSP90B1, crucial for TLR-dependent immune responses ^27,28^. Our findings demonstrate that CNPY3 is a key gene associated with intrinsic immunity, predominantly expressed in immune cells. While previous research has largely focused on the role of CNPY3 in early infantile epileptic encephalopathy ^29,30^, its potential involvement in flavivirus infections remained unexplored.

Our study demonstrates that flavivirus infection significantly downregulates the expression of CNPY3, a process mediated by both the positive and negative RNA strands of the virus. This downregulation occurs through the disruption of the interaction between HuR and CNPY3 mRNA, which destabilizes the CNPY3 mRNA and reduces its expression. Previous studies have shown that HuR is involved in post-transcriptional regulation of various mRNAs and plays a crucial role in stabilizing transcripts that are essential for immune responses^31–33^. Our findings align with these studies, as we reveal that upregulating CNPY3 can inhibit flavivirus replication, whereas downregulating CNPY3 impairs the host immune response and enhances viral infection. Collectively, these results provide strong evidence that CNPY3 is a critical factor in the host’s immune defense against flavivirus infections. The virus appears to exploit the HuR protein to destabilize CNPY3 mRNA, thereby suppressing its expression and evading the host immune response. This mechanism underscores the complex strategies employed by flaviviruses to modulate host cell machinery for their benefit.

What particularly intrigued us was the involvement of the negative strand RNA of flaviviruses in this process. It is well-established that the negative strand RNA acts as an intermediate during viral replication. However, its role in immune evasion remains poorly understood. Several studies have highlighted interactions between host cellular proteins and the negative strand RNA of flaviviruses, such as the binding of host proteins to the 3’ untranslated region of DENV-2 ^34^ and the 3’ stem-loop RNA of West Nile virus ^35^. However, the implications of these interactions for viral replication and immune escape mechanisms have not been extensively investigated. Our study provides novel evidence that flavivirus negative strand RNA actively participates in immune evasion, revealing a new layer of complexity in the interaction between host immune defenses and viral infection.

Upregulation of CNPY3 effectively inhibited replication of both DENV and ZIKV, suggesting that CNPY3 holds potential as a therapeutic target for flavivirus diseases. mRNA therapy presents several advantages, including ease of synthesis and modify nucleotide to encode specific proteins, high potency, rapid development, low-cost manufacturing, and safe administration. These benefits facilitate the swift development of mRNA-based treatments for various diseases^36–38^. While mRNA vaccines delivering viral antigens have received significant attention, there are fewer reports on mRNA therapeutics encoding antiviral proteins for treating existing infections ^39–41^. Our study demonstrates the potential of mRNA therapy by developing LNP-CNPY3 as an antiviral treatment for DENV and ZIKV infections in a mouse model. As anticipated, this approach significantly reduced encephalitis symptoms and improved survival rates. Notably, this represents a novel application of mRNA therapy, extending beyond vaccine development to include post-infection treatment. However, the intracranial injection method used in our study is invasive and potentially damaging to the brain. Future research should explore less invasive delivery methods for CNPY3 mRNA to enhance its therapeutic potential.

In summary, our study identifies a new therapeutic target against flaviviruses and evaluates its potential as an mRNA-based drug therapy in animal models. We also uncovered a novel mechanism by which flavivirus negative strand RNA contributes to immune evasion. This pioneering research not only elucidates the complex interaction between CNPY3 and flavivirus pathogenesis but also paves the way for innovative therapeutic strategies leveraging CNPY3 modulation. Our findings have significant implications for understanding host-virus interactions and offer promising new avenues for targeted antiviral interventions.

## Materials and Methods

### Primary Analysis

The datasets related to Dengue Virus (DENV) and Zika Virus (ZIKV) analyzed in this study were obtained from the Gene Expression Omnibus (GEO). These datasets include microarray data of DENV-infected dendritic cells (GSE58278), transcriptome data of DENV patients (GSE51808), transcriptome data of ZIKV exposure in human cortical neural progenitors (GSE80434). Differential expression analysis of the datasets was performed using the NCBI online tool GEO2R^42^. Genes with p-value ≤ 0.05 and Log2 fold change (Log2FC) ≥ 1 between the infected and control groups were considered as differentially expressed genes (DEGs). Venn analysis was conducted using online tools to generate custom Venn diagrams (http://bioinformatics.psb.ugent.be/webtools/Venn/). The Kyoto Encyclopedia of Genes and Genomes (KEGG) pathways and Gene Ontology (GO) categories associated with the up-regulated and down-regulated DEGs were visualized using Metascape (https://metascape.org/gp/index.html#/main/step1)^43^.

### Cell Culture and flavivirus infection

The strains of DENV-1 (THD1-0102-01), DENV-2 (New Guinea), DENV-3 (80-2), DENV-4 (GD07-78) are provided by the Center for Disease Control and Prevention of Guangzhou Military Command, China. ZIKV strains FSS13025 kindly provided by Cheng-Feng Qin (Professor of Virology, Beijing Institute of Microbiology and Epidemiology). For Vero, HEK 293T and A549 cells culture, they all used the DMEM (Gibco, Life Technologies, USA) supplemented with 10% FBS. For THP-1 and U937 cells, they were cultured in RPMI-1640 medium (Invitrogen-Gibco, USA) supplemented with 10% fetal bovine serum (FBS). The culture medium was aspirated, and the cells were washed twice with serum-free medium. Subsequently, flavivirus at a multiplicity of infection (MOI) of 2 was added to the cells and incubated for 2 hours. Then the virus was removed, and the cells were further cultured in 24-well plates for different time points to further analysis.

### Animal Infection

Animal infection was conducted following the previously described method^44^. DENV-2 was injected into 3-day-old suckling mice (female BALB/c pregnant mice were obtained from the Experimental Animal Center of the Third Military Medical University). For intracranial infection, 3-day-old BALB/c mice were intracranially infected with 10 μL of DENVs or ZIKV or isopycnic PBS as the control group. Then expression of CNPY3 in whole blood was then measured in the mice at different time points post-infection.

For intranasal infection, DENV-2 which isolated from the brain tissue of suckling mice was administered as nasal drops to 3-day-old suckling mice in a volume of 3 μL. The isopycnic supernatant from the brain tissue of healthy mice was applied as the control group. Subsequently, RNA was extracted from the brain and blood samples at different time points after infection.

### Quantitative real-time PCR (RT-qPCR)

RNA was extracted from the collected cells and mouse tissues using the Tissue/Cell RNA Rapid Extraction Kit (BOER, China) following the manufacturer’s instructions. The extracted RNA was then subjected to reverse transcription using the PrimeScriptTM RT Reagent Kit (TaKaRa, Japan). Quantitative real-time PCR was performed using TB Green® Premix Ex Taq™ II (TaKaRa, Japan) and the LightCycler® 96 System (Roche, USA). The experimental data of PCR reaction was analyzed for abnormality by checking the dissolution curve and amplification curve. The expression level of the target gene was calculated with the −ΔΔct method. The primer sequences used for target gene detection are provided in Data files S1.

### Transient transfection

*In-vitro* transient transfection of CNPY3 was performed in HEK 293T cells or A549 cells using Lipofectamine® 3000 reagent (Invitrogen, USA) according to the manufacturer’s instructions. Briefly, cells were transfected with equal concentrations of pCDNA3.1-CNPY3 constructs or vector controls in serum-free DMEM medium (Gibco, Life Technologies, USA). After 24 hours of incubation, complete medium was added to the cells. Transient transfection of CNPY3-targeted siRNA into THP-1 cells was carried out using Zeta Life Advanced DNA/RNA transfection reagent (Zeta Life, USA) as instructed by the kit protocol. The cells were inoculated at a density of 3×10^5^ cells/mL in 6-well cell culture plates with 2 mL in each well. The synthesized negative control siRNA (NC) and CNPY3 siRNA were dissolved in nuclease-free water to a final concentration of 20 uM. The volume of siRNA and transfection reagent was mixed at a 1:1 ratio and incubated at room temperature for 15 min. Functional analyses were conducted after culturing 36 h. The sequences of siRNAs used are provided in Data files S1.

### Detection of DENV Envelope (E), NS3 and NS5 genes expression

HEK 293T and THP-1 cells were transiently transfected with pCDNA3.1-CNPY3 or pcDNA3.1 empty plasmid or siRNA for 36 h. Subsequently, the transfected cells were infected with DENV-2 at MOI of 2 for 18 h. RNA was then extracted from these cells and reverse transcribed into cDNA for measuring the relative expression levels of DENV-2 E, NS3, and NS5 genes. The primer sequences used for detecting the E, NS3, and NS5 genes are provided in Data files S1.

### Copy number detection of virus in cells supernatant

Viral RNA from the flavivirus infected cells supernatant was extracted using the MiniBEST Viral RNA/DNA Extraction Kit (Takara, Japan) and eluted with 30 μL RNase-free water. Then 7 μL of the RNA was used for reverse transcription by the PrimeScriptTM RT Reagent Kit (TaKaRa, Japan). To obtain a total of 20 μL cDNA. For quantitative real-time PCR (qRT-PCR), 1 μL cDNA was taken as a template. The RNA copies per 200 μL of each sample were calculated from the qRT-PCR Cq values using a standard curve. The primers used for qPCR are provided in Data files S1.

### Plaque Assay

Vero cells were seeded in 24-well plates and grown to form monolayers. Virus samples were added to the Vero cells and incubated at 37°C for 2 h. The inoculum was then removed, and the monolayers were overlaid with 1.5 mL of DMEM containing 1.2% methylcellulose. The cells were further incubated at 37°C for 6 d and then fixed using 4% paraformaldehyde. Finally, the plaques were stained with 1% crystal violet and counted.

### Cell immunofluorescence analysis

HEK 293T or A549 cells were seeded on 24-well tissue culture plates and transfected with pcDNA3.1-CNPY3 or pCDNA3.1 empty plasmid. The transfected cells were then infected with DENV-2 or ZIKV at MOI of 2. Subsequently, the cells were fixed with 4% paraformaldehyde solution at room temperature for 15 min and treated with 0.5% Triton X-100 solution (preheated to 37°C). The cells were incubated with anti-dengue Virus 1+2+3+4 antibody (Abcam, England) or anti-flavivirus group antibody D1-4G2-4-15 (4G2) (1:200, Genetex, USA) at 4°C overnight. After washing the cells with PBS twice, they were stained with an Alexa Fluor® 488-labeled sheep anti-rabbit IgG (H+L) secondary antibody (Abcam, England) at 37°C for 1 h. The cells were scanned using a fluorescence microscope (Olympus, Southborough, MA, USA), and the number of virus-infected cells in a specific region was counted using Image J software ^45^.

### Western blot analysis

The collected samples were washed twice with PBS to remove residual serum and then lysed on ice for 20 min. The protein concentration was determined using a BCA protein concentration determination kit (Beyotime, China). The samples were denatured at 100°C for 10 min. The proteins were separated by 12% SDS-PAGE and transferred to a polyvinylidene fluoride membrane (Beyotime, China). The membrane was blocked with 2% BSA for 1 h and incubated with antibodies against CNPY3 (Sino Biological, China), anti-DENV E/Envelope protein (Sino Biological, China), and β-actin antibody (Beyotime, China) at 4°C overnight. After the membrane washed with PBS containing 0.1% tween (PBST) three times, horseradish peroxidase (HRP)-labeled secondary antibody was added and incubated at room temperature for 1 h. Blots were developed with chemiluminescence substrate ECL (Thermo Scientific, USA) and visualized with a Bio-Rad imaging system.

### Preparation of lipid nanoparticles (LNP)

*Cnpy3* mRNA was synthesized using the T7 High Yield RNA Transcription kit (Novoprotein, China) with linearized plasmids (puc57-Cnpy3, 5′UTR-Cnpy3 ORF-3′UTR). Sequences showed in Data files S1. Cap1 was added to the synthesized RNA using the Cap 1 Capping System (Novoprotein, China), and a poly(A) tail was generated using E. coli Poly(A) Polymerase (Novoprotein, China).

The LNP-mRNA formulations were prepared as previously described^46^. SM102 (AVT, China), DSPC (AVT, China), cholesterol (AVT, China), and DMG-PEG2000 (AVT, China) were solubilized in ethanol at a molar ratio of 50:10:38.5:1.5. After equilibrating at room temperature for 2 min, the lipid mixture and mRNA dissolved in 50 mM citrate buffer were added into a microfluidic chip. The lipid phase was mixed with the aqueous phase (mRNA) at a flow rate of 1:3 at room temperature, forming vesicles of approximately 100 nm in diameter. The mRNA, solubilized in 50 mM citrate buffer, was mixed with lipid at a ratio of 6:1, resulting in a final ethanol concentration of 30% (v/v) and a lipid concentration of 6.1 mg/mL. The LNP-encapsulated mRNA samples were dialyzed against PBS (pH 7.4) in dialysis bags (Viskase, USA) for 24 h and then stored at 4°C until use. The encapsulation efficiency was measured using the Quant-iT RiboGreen RNA Assay Kit (Invitrogen, USA) with a Varioskan lux (Thermo Scientific, USA) instrument.

### Characterization of Lipid-*Cnpy3* mRNA nanoparticle

The hydrodynamic diameters of the Lipid-Cnpy3 mRNA nanoparticles were measured at 25°C using a Zetasizer Nano ZS90 instrument (Malvern Panalytical Ltd, U.K.). The morphology of the nanoparticles was observed using a transmission electron microscope (TEM), specifically the Tecnai G2 F20 S-TWIN (FEI Company, USA).

### Measurement of type I interferon concentrations by ELISA

Mouse Ifn-α and Ifn-β, as well as human IFN-β, were quantified using enzyme-linked immunosorbent assay kits from MEIMIAN (China) and MultiSciences (China), following the manufacturer’s instructions.

### RNA-Sequencing

Total RNA was extracted from brain tissues using the TRIzol® Reagent (Invitrogen, USA) following the manufacturer’s instructions. Genomic DNA was removed using DNase I (TaKaRa, Japan). RNA-seq libraries were prepared using the TruSeqTM RNA sample preparation Kit from Illumina (San Diego, CA) with 1 μg of total RNA. The libraries were quantified using TBS380, and paired-end RNA-seq sequencing was performed using the Illumina HiSeq xten/NovaSeq 6000 sequencer with a read length of 2 ×150bp. The reads were trimmed, and quality controlled using SeqPrep and Sickle with default parameters. The clean reads were aligned to the reference genome using HISAT2 version 2.1.0^47^. The mapped reads of each sample were assembled using StringTie version 2.1.2^48^ in a reference-based approach. The expression levels were calculated using the transcripts per million reads (TPM) method, and gene abundances were quantified using RSEM version 1.3.3. Data analysis was performed using R version 3.3.3 and DESeq2 version 1.24.0. Differential gene expression analysis was conducted based on a Q value ≤ 0.05 and |log2FC| > 1. The data were analyzed on the online platform of Majorbio Cloud Platform (www.majorbio.com)^49^.

### Brain Histology

Paraffin sections of brain tissues were dewaxed with xylene and hydrated with ethyl alcohol. For hematoxylin and eosin stain, the sections were stained with hematoxylin for 3 min and eosin for 5 min. For Nissl staining, duplicate sections were stained with 0.75% cresyl violet, dehydrated through graded alcohols. Finally, all sections were digitalized using a Zeiss Axio Scan.Z1 slide scanner.

### Brain Immunofluorescence

Brain immunofluorescence was followed the method described previously with modified^44^.

Paraffin-embedded brain sections were dewaxed and treated with a 3% hydrogen peroxide solution for 25 min. The sections were then boiled in 0.01 M citrate buffer (pH 6.0) for 10 min and incubated with normal goat serum (Abcam, Cambridge, UK) for 2 h at 37 °C. Next, the sections were incubated with primary antibodies, including Anti-Flavivirus NS1 at 1:500 (Abcam, UK), Anti-IBA1 at 1:200 (WAKO, Japan), Anti-Flavivirus 4G2 (1:200, Genetex, USA), Anti-NeuN (Abcam, UK) or Anti-HuR at 1:100 (Bioworld Technology, Inc. China) overnight at 4 °C. Subsequently, the sections were incubated with FITC-conjugated or CY3- conjugated secondary antibodies (Servicebio, Wuhan, China) for 2 h. Finally, the sections were stained with DAPI (Abcam, UK) for 4 min. Representative sections were scanned using a Pannoramic DESK digital slice scanner (3D HISTECH, Hungary) with the Pannoramic Scanner software.

### Fluorescence in situ hybridization (FISH) and immunofluorescence (IF) analysis

RAW264.7 cells were grown on glass coverslips, after DENV-2 infected or uninfected for 48 h (MOI = 1), fixed in 4% paraformaldehyde for 15 min. Fluorescence in Situ Hybridization Kit for RNA (Beyotime, China) was used for FISH analysis. Briefly, after prehybridization at 55 °C for 20 min, coverslips were hybridized with Cy3-labelled CNPY3 probes (APExBIO, USA) and FITC-labelled DENV-2 probes (Yemai biotechnology, China) at 55 °C overnight, and then washed with Washing Buffer. The sequence of the probe used is listed in Data files S1. Then, the coverslips were incubated with an antibody specific for HuR (1:200, Bioworld Technology, USA) at 37◦C for 2 h and washed with PBS. Subsequently, Alexa Fluor 647 goat anti-rabbit IgG (1:800, Abcam, England) was added to the coverslips at 37◦C for 1 h and washed with PBS. Finally, the nuclei were stained with DAPI (Beyotime, China), and the fluorescence signal was visualized under a confocal microscopy (ZEISS, LSM880, Germany).

### Flow cytometry

A549 cells were seeded on 6-well tissue culture plates and transfected with pcDNA3.1- CNPY3 or pCDNA3.1 empty plasmid for 24 h. The transfected cells were then infected with ZIKV at an MOI of 2 for 48 h. Subsequently, cells were fixed and permeabilized using Intracellular Fixation&Permeabilization Buffer Set (Invitrogen, USA). Cells were incubated with an anti-flavivirus group antibody D1-4G2-4-15 (4G2) (1:200, Genetex, USA) and FITC-labelled goat anti-mouse IgG (1:800,Abcam, England) secondary antibody at 4°C for 1 h. Finally, cells were analyzed on a BD flow cytometer and the data were processed using FlowJo software.

### RNA-Pull-down and RNA immunoprecipitation (RIP)

To confirm that CNPY3 and DENV-2 RNA sequences can bind to the HuR protein, we used an F2-RNA-Pull-down kit (FitGene Biotechnology Co., Ltd., China), following the manufacturer’s instructions. The F2 tag is a short 16-nucleotide RNA sequence (GGCGCTGACAAAGCGC) fused to the target RNA sequence. The 3’ UTR of DENV-2 and its complementary negative strand were synthesized and inserted into the PUC57 vector (GenScript Biotech Corporation, China). These plasmids, along with the pcDNA3.1-CNPY3 plasmid, were used as templates to generate the target RNA molecules via PCR. The PCR used an upstream primer containing the T7 promoter and a downstream primer containing the F2 tag. The PCR products were used as templates for RNA transcription using T7 RNA polymerase (Novoprotein, China). Then the target RNA was denatured at 95°C for 3 minutes before being bound to magnetic beads.

To obtain enough HuR protein, we transfected 293T cells with the pcDNA3.1-HuR plasmid (Boer, China) and harvested total protein 36 hours post-transfection. Afterward, total protein was added to the magnetic beads which bounded target RNA and incubated at 4°C for 2 hours. The magnetic beads were washed twice to remove unbound proteins, and the bound proteins were analyzed by Western blot using an anti-HuR antibody (Bioworld, China).

To assess the association of endogenous HuR with the DENV-2 UTR and CNPY3 mRNA, RNA immunoprecipitation (RIP) was performed using a RIP kit (BersinBio, China). The DENV-2 negative strand RNA was transfected into 293T cells, and a total of 1×10^7 cells were collected per sample. The lysates were divided into two portions: 0.8 mL for immunoprecipitation (IP) and 0.1 mL as input. For the IP samples, 5 μg of IP antibody (IgG or anti-HuR) was added and incubated overnight at 4°C on a vertical mixer. The RNA from the IP samples was extracted using a phenol-chloroform-isoamyl alcohol mixture, and qPCR was used to measure the levels of CNPY3 and DENV-2 3’ UTR negative strand RNA.

The primers used for RNA pull-down and RIP are listed in Data files S1.

### Ethical statement

All animal experiments were conducted following the guidelines of the Chinese Regulations of Laboratory Animals. The protocol was approved by the Animal Care and Use Committee of the Army Medical University and followed the National Institutes of Health Guide for the Care and Use of Laboratory Animals.

### Statistical analysis

All statistical analyses were performed using GraphPad Prism v.7 and v.9 software. Two-tailed unpaired Student’s *t*-test was used for comparison of between-group data. One-way analysis of variance (ANOVA) followed by Dunnett’s test was used for multiple comparison. Survival rates were statistically analyzed using the log - rank (Mantel–Cox) test.

## Acknowledgments

This work is supported by the Natural Science Foundation Project of Chongqing grant cstc2020jcyj-bshX0116 (XYD). We would like to express our sincere gratitude to Professor Yan Pei for his valuable insights on this study. We also extend our thanks to Dr. Chiara Lincetto for her assistance with language editing.

## Author contributions

Conceptualization, XYD, JTL

Methodology, XYD, JTL

Software, XYD Validation, XYD

Cell experiment and animal experiment, XYD, JXH, YXZ, JZ, JT, WXQ

RIP and RNA-pull-down, XYD, XZC

mRNA delivery platform, SG

Virus culture, MYQ, DH, MCL

Data curation, XYD

Writing—original draft: XYD, HJX

Writing—review & editing: XYD, HJX, SG, JTL

## Data and materials availability

The RNA seq data has been deposited to the GEO database with the accession code GSE234119 (https://www.ncbi.nlm.nih.gov/geo/query/acc.cgi?acc=GSE234119).

## Supplementary Materials

### Supplementary Figures

**Fig S1.**
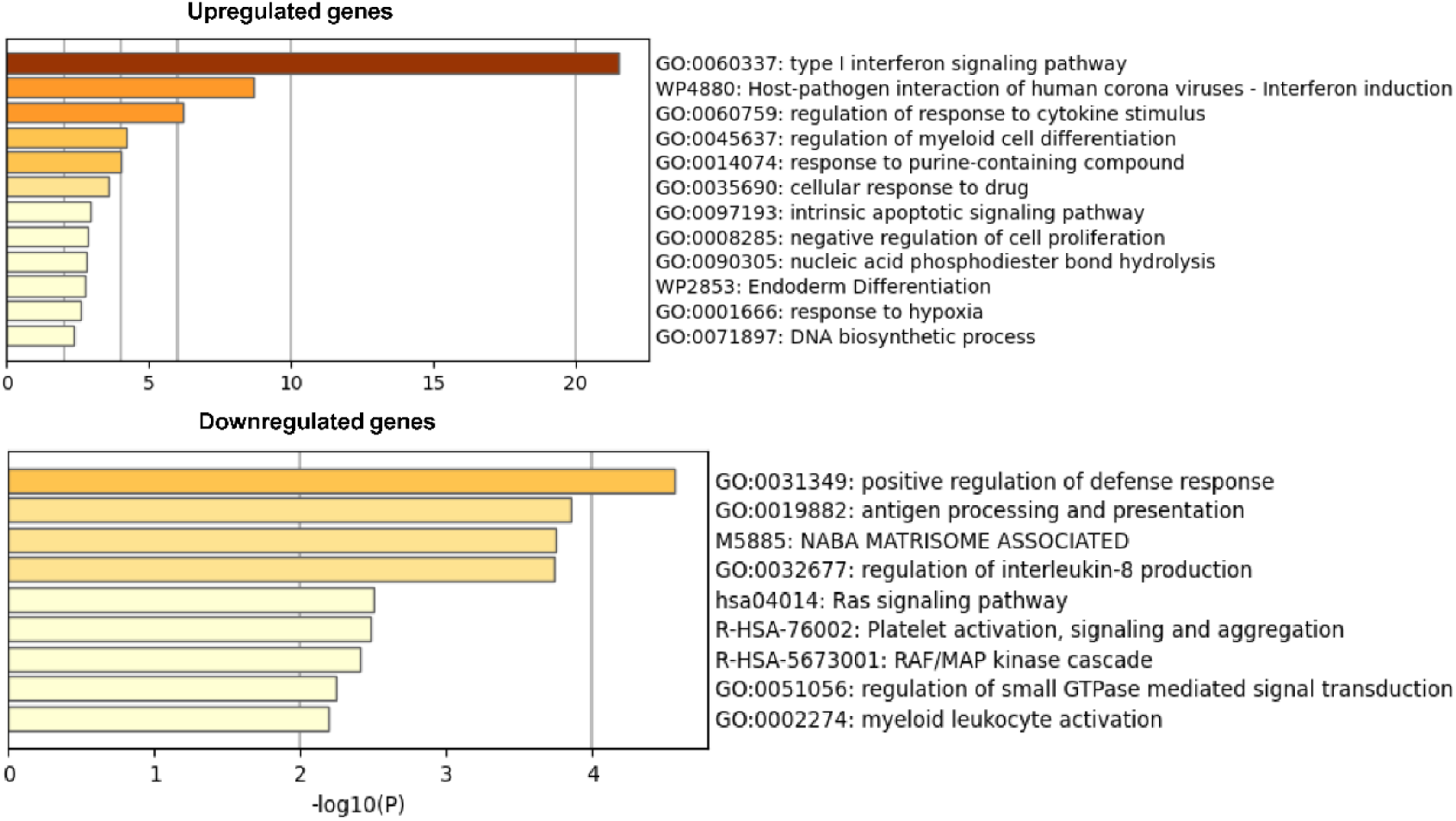
Functional enrichment analysis of DEGs. Heatmap shows top gene ontology (GO) biological processes. Discrete color scale represents statistical significance.

**Fig S2.**
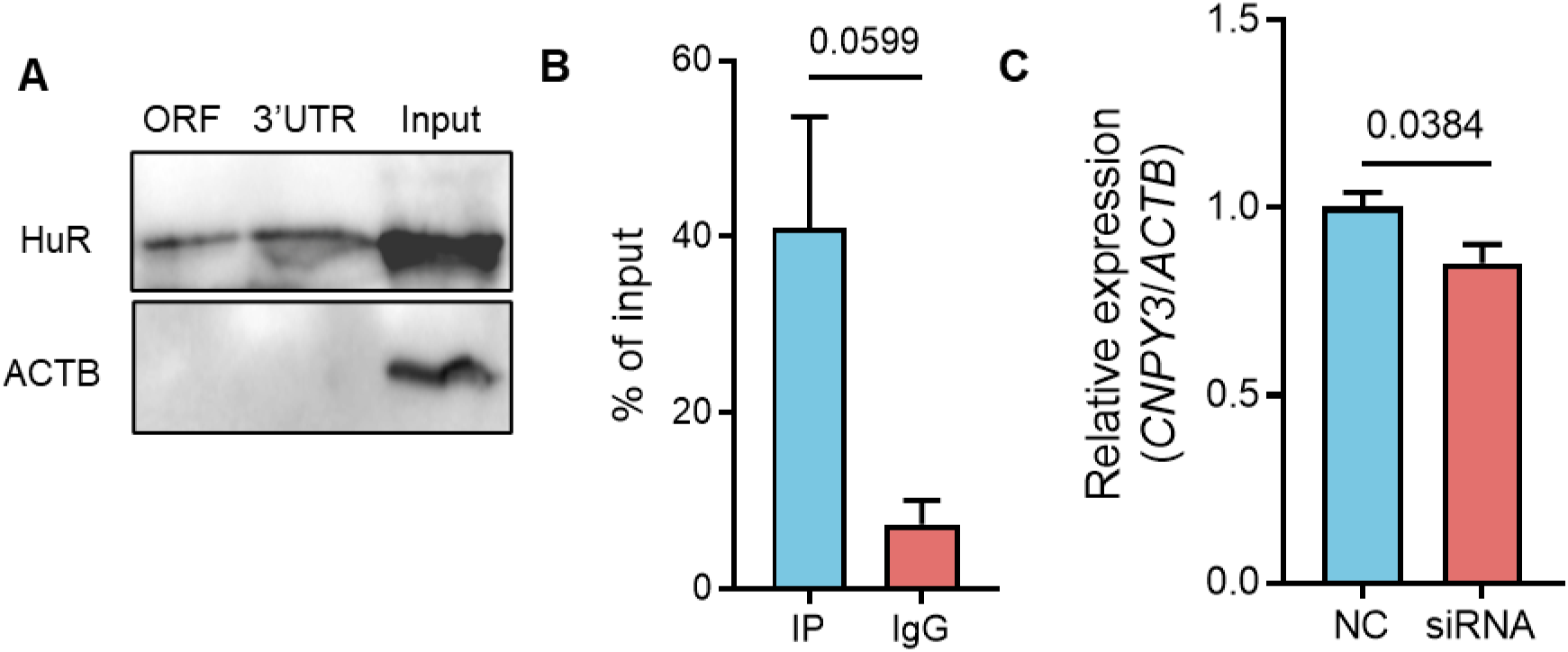
HuR protein binds to CNPY3 mRNA and regulates CNPY3 expression. (A) RNA-pulldown was used to detect the interaction between CNPY3 mRNA and RNA binding protein HuR. ORF, CNPY3 Open Reading Frame; 3′UTR, 3′ untranslated regions; Input, positive control. (B) RIP analysis was used to measure the association of HuR with CNPY3 mRNA 3′UTR by using either anti-HuR antibody or control IgG. (C) CNPY3 expression in U937 cells treated with CNPY3-siRNA or Negative Control (NC)-siRNA. Results are presented as mean ±SEM (n = 3). P values were determined by an unpaired two-tailed *t*-test.

**Fig S3.**
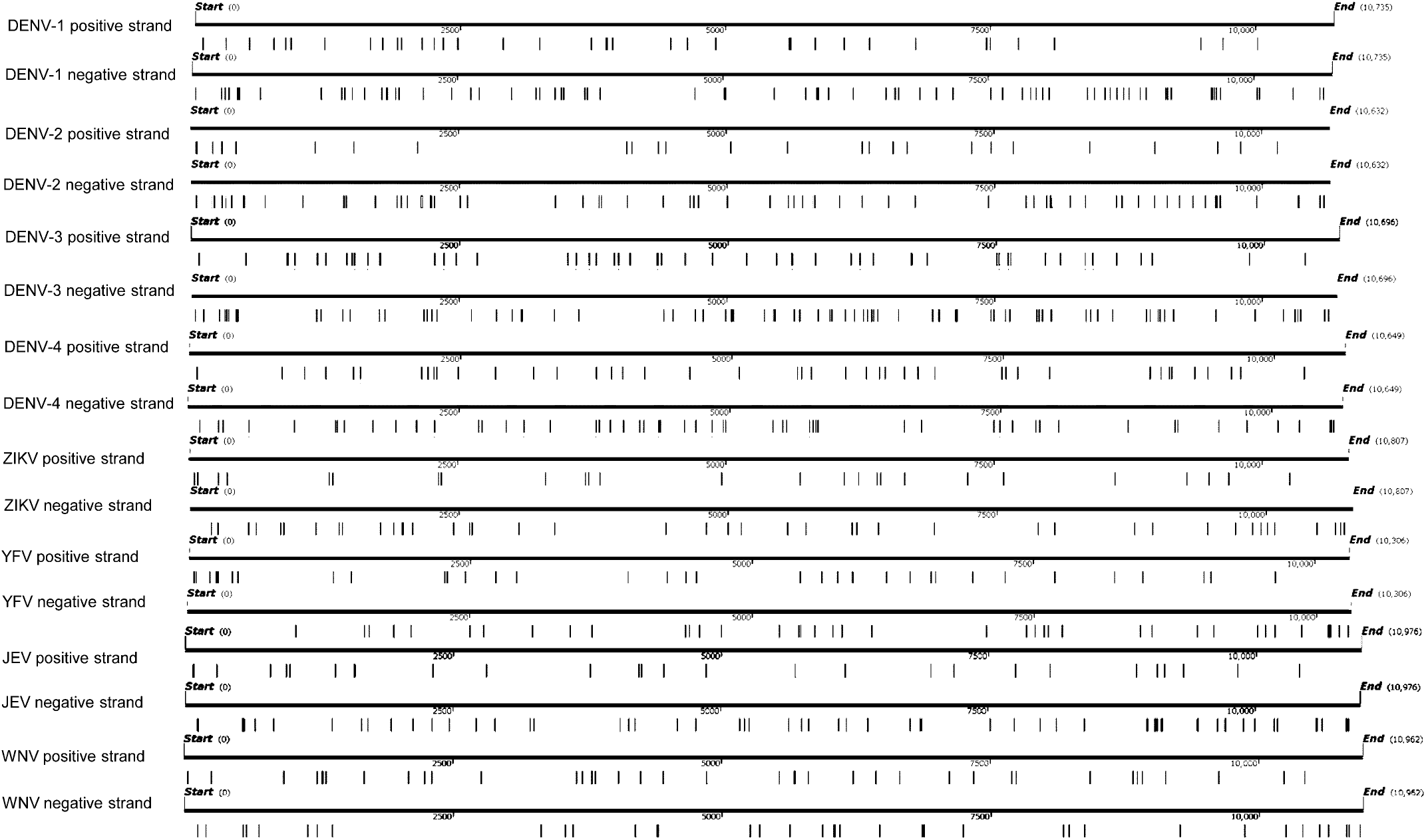
The location and the amounts of AU-rich elements (AREs) in Flavivirus positive and negative strand. DENV, Dengue virus; ZIKV, Zika virus; YFV, Yellow Fever virus; WNV, West Nile virus; JEV, Japanese Encephalitis. The black vertical lines represent ARE.

**Fig S4.**
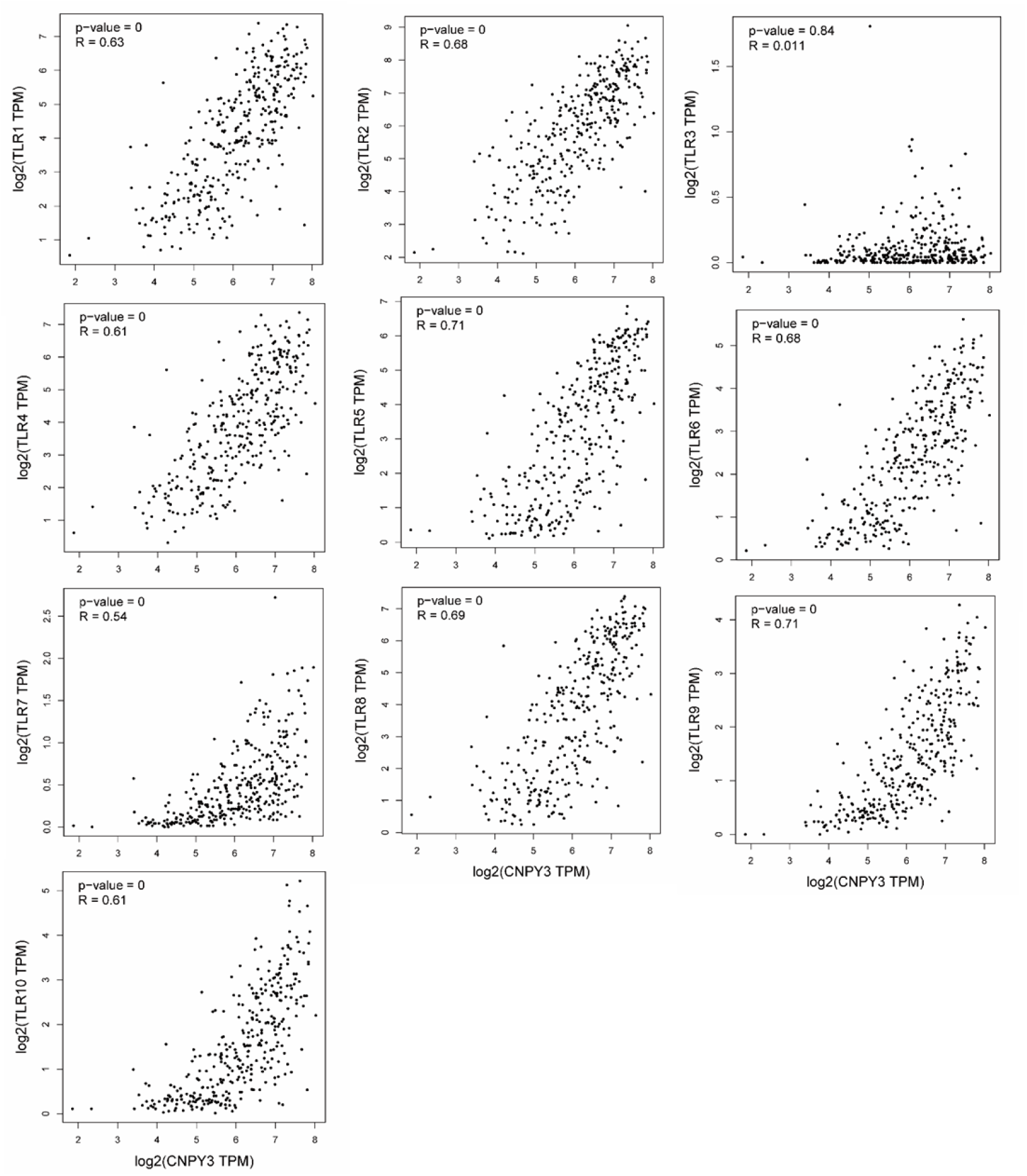
Correlation analysis of CNPY3 and Toll-like receptor expression in healthy blood based on GEPIA.

**Fig S5.**
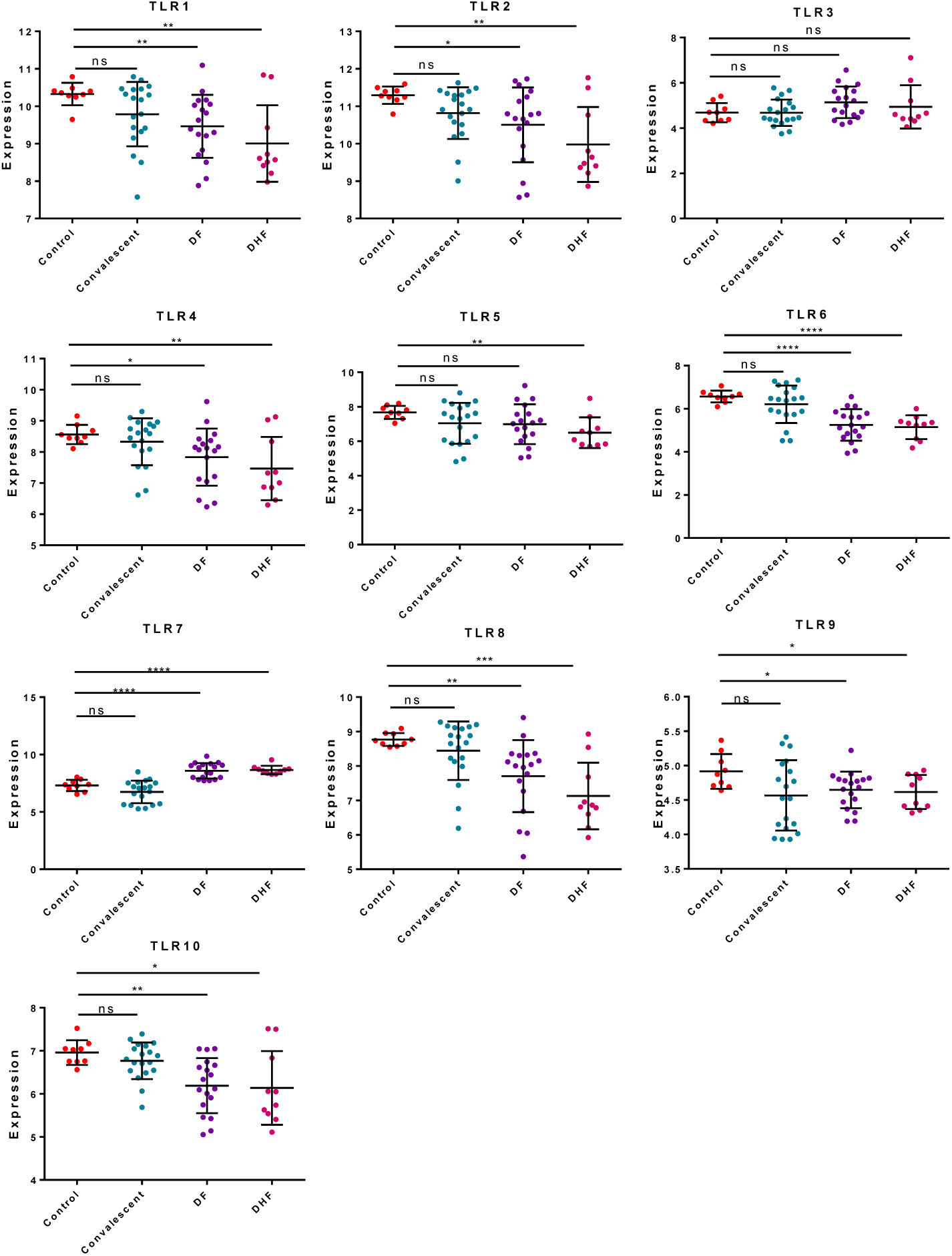
Correlation analysis of CNPY3 and Toll-like receptor expression in DENV patients’ blood based on GSE51808.

**Fig S6.**
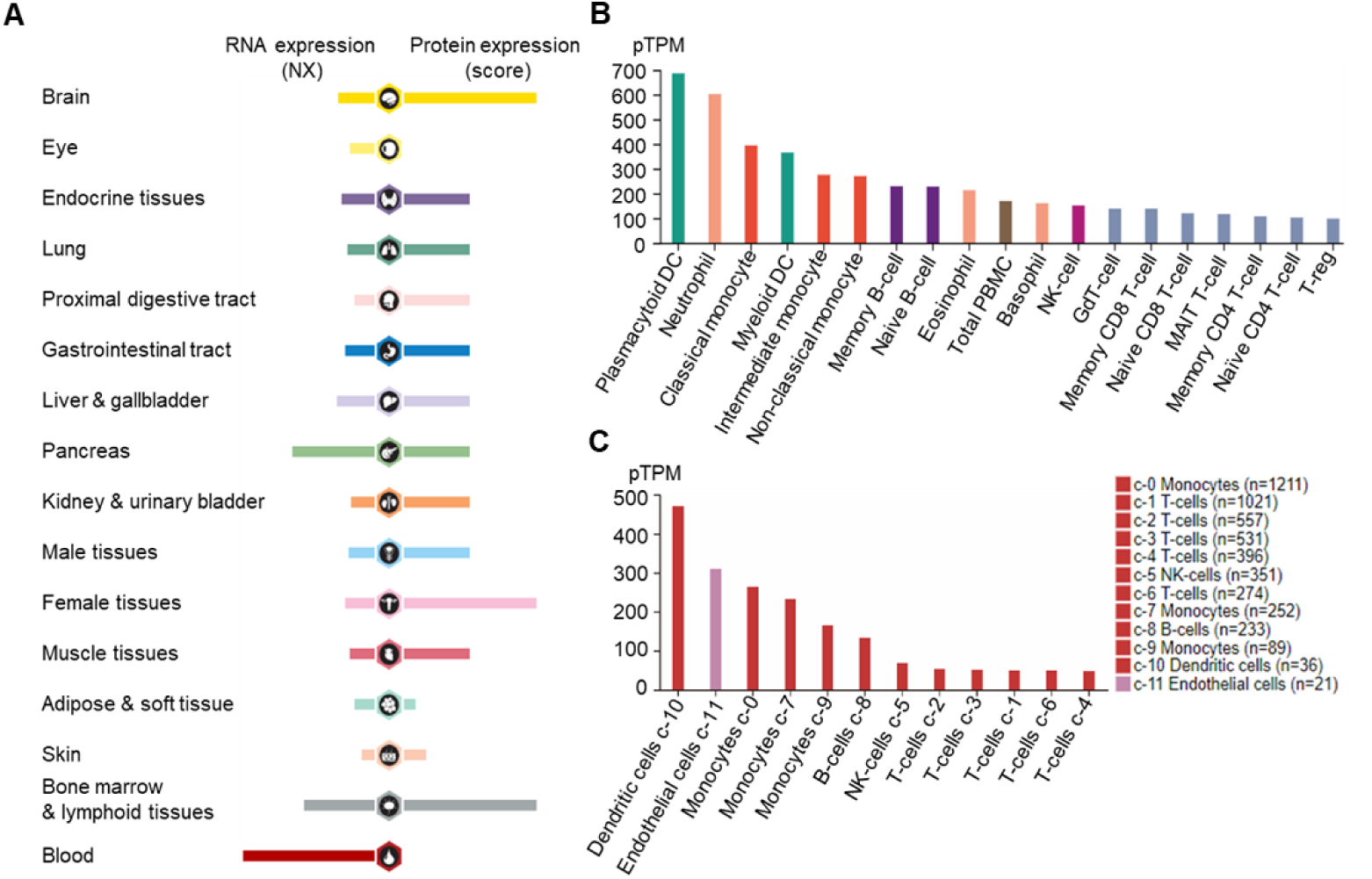
Expression specificity analysis of CNPY3 in human tissues and blood cells using Human Protein Atlas database. (A) CNPY3 expression analysis at mRNA and protein levels in human tissues. (B) CNPY3 expression in human blood cells was analyzed b HPA scaled dataset. (C) CNPY3 expression in blood on a single cell level. Single cell data includes scRNA-seq data from clusters of various peripheral blood mononuclear cell types (PBMC) and endothelial cells.

**Fig S7.**
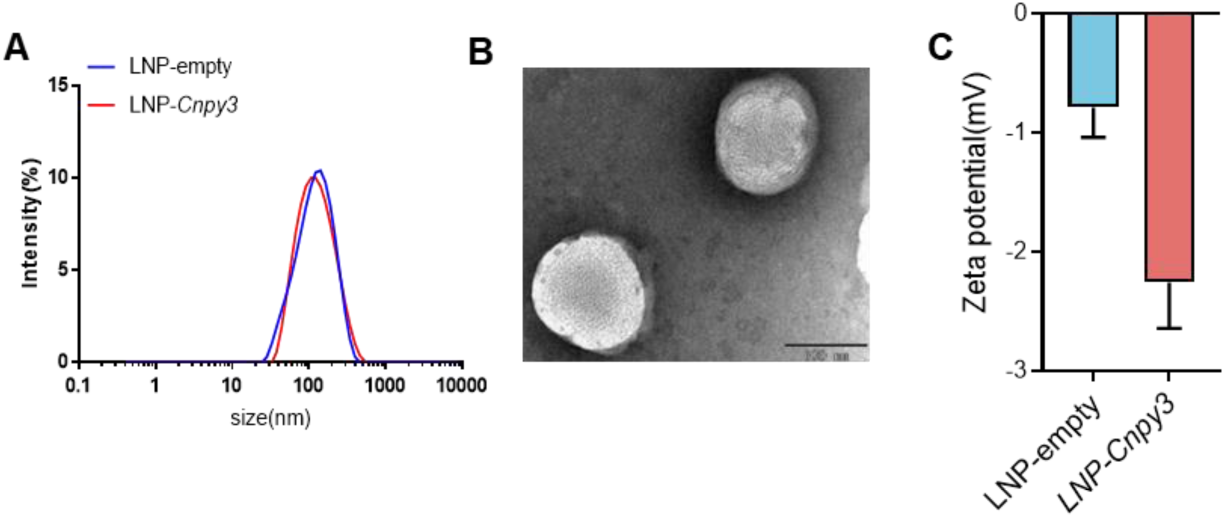
Nanoparticle properties and characterization. (A) Particle size detection of LNP-Cnpy3 and LNP-empty using DLS laser particle size analyzer (B)TEM image of LNP-Cnpy3, scale bar = 100 nm. (C) ZETA potential detection of LNP-mRNA using DLS laser particle size analyzer.

**Fig S8.**
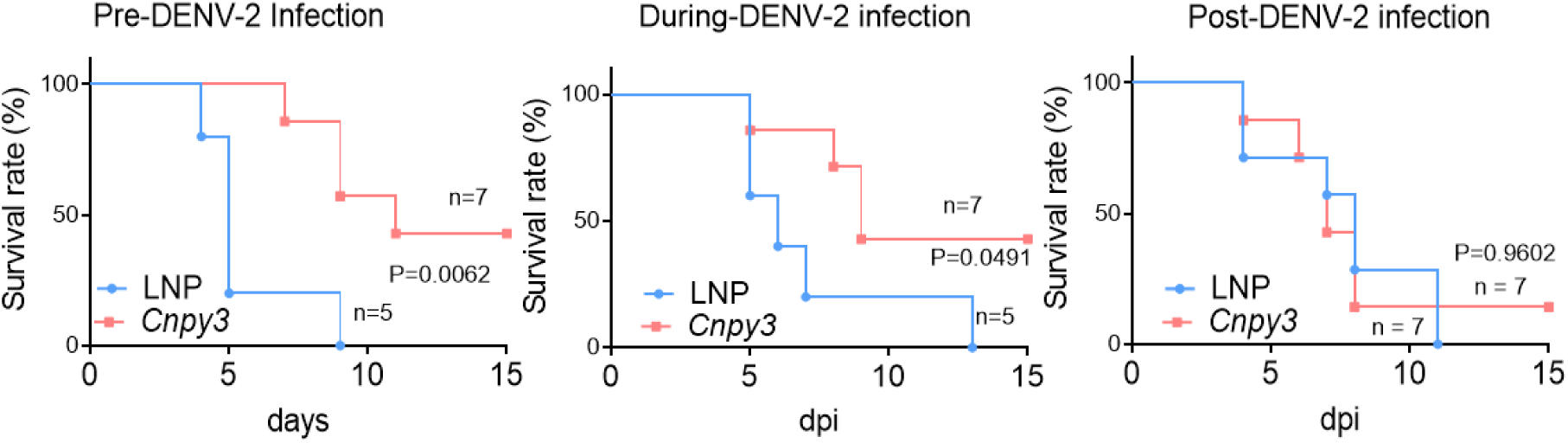
Efficacy of early and delayed administration of LNP-Cnpy3. DENV-2-infected mice were administered LNP-Cnpy3 or LNP on -1-0-dpi (Early infected), 0-1-dpi (During infected), 3-4 dpi (Post infected). Survival rates were statistically analyzed using the log-rank (Mantel–Cox) test.

**Fig S9.**
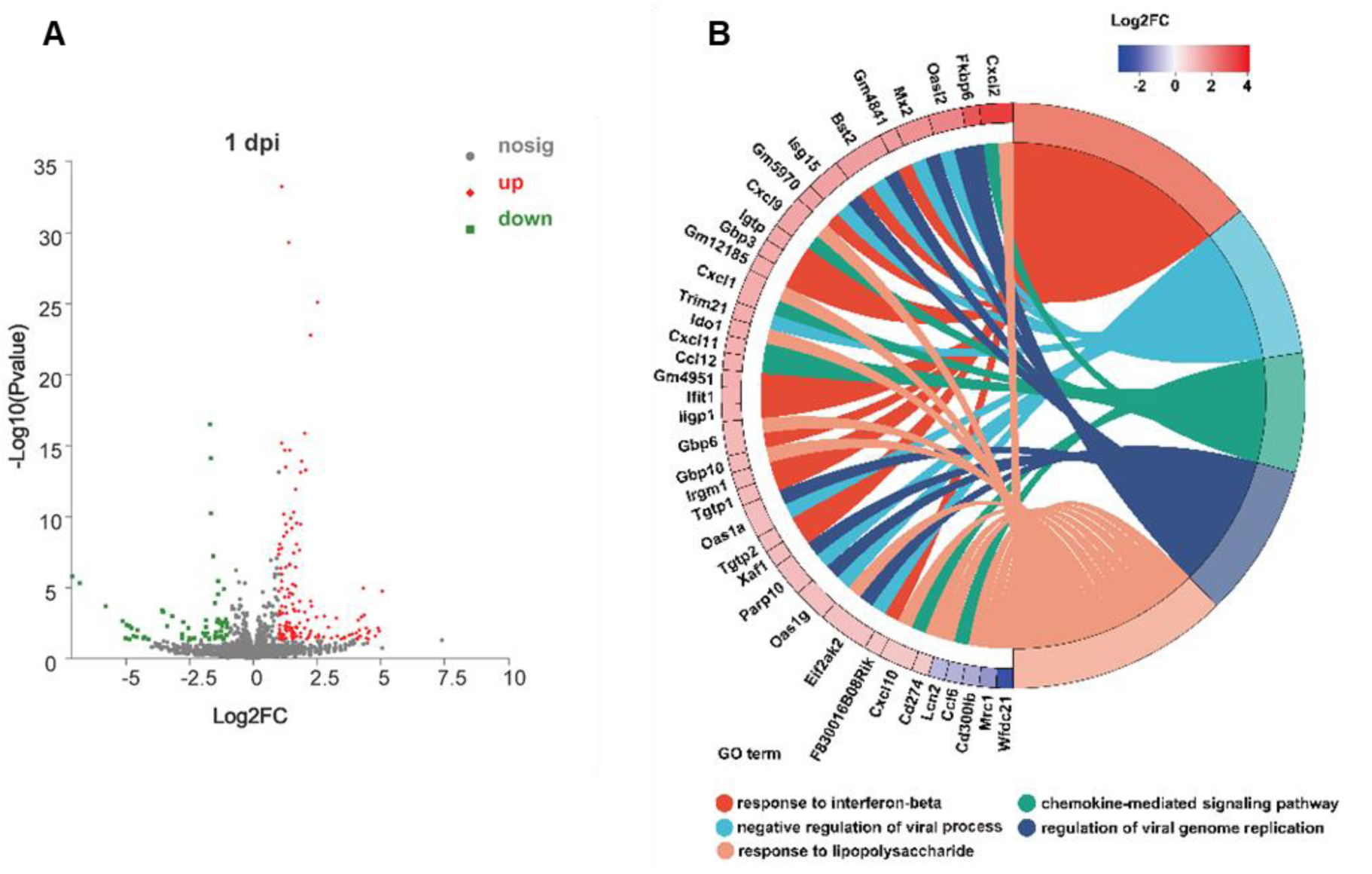
Transcriptomics analysis in early stage flavivirus infection. (A) Volcano plot based on differentially expressed genes in samples. (B) Ontology (GO) enrichment analysis. RNA-Seq analysis of brains from 3-day-old BALB/c mice brains infected with DENV-2 and treated with either LNP-*Cnpy3* or LNP alone on 1-day-post-infection (dpi). (A) Ontology (GO) enrichment analysis of DEGs in brain tissues from Lipid-Cnpy3 mRNA or Lipid-empty-treated DENV-2 infected pups at 1 dpi. Chords represent a detailed relationship between the expression levels of DEGs (left semicircle perimeter) and their enriched GO pathways (right semicircle perimeter). For each gene, the expression value (log2 FC) of DEGs in brain tissue is shown by colored rectangles. (B) Genes of related to response to interferon-beta are presented by heatmap based on gene expression value. RNA-Seq analysis of brains from Lipid-Cnpy3 mRNA or Lipid-empty-treated DENV-2 infected pups at 1 dpi (n=3 pups respectively). log2 FC are color coded as indicated.

**Fig S10.**
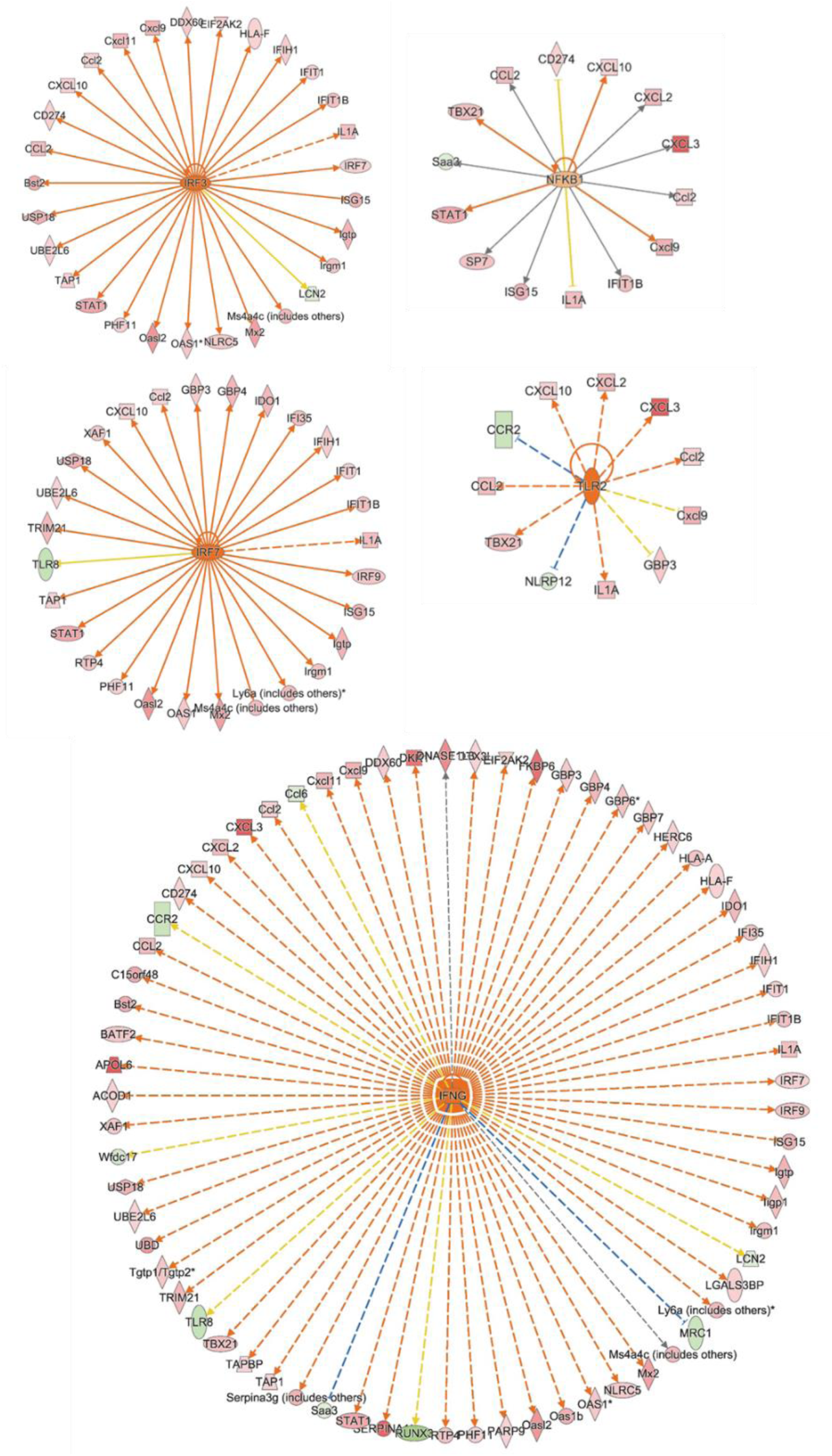
IPA identified IRF3, IRF7, IFNG, NFKB1 and TLR2 as upstream regulators. Orange for up- regulation, green for suppression.

**Fig S11.**
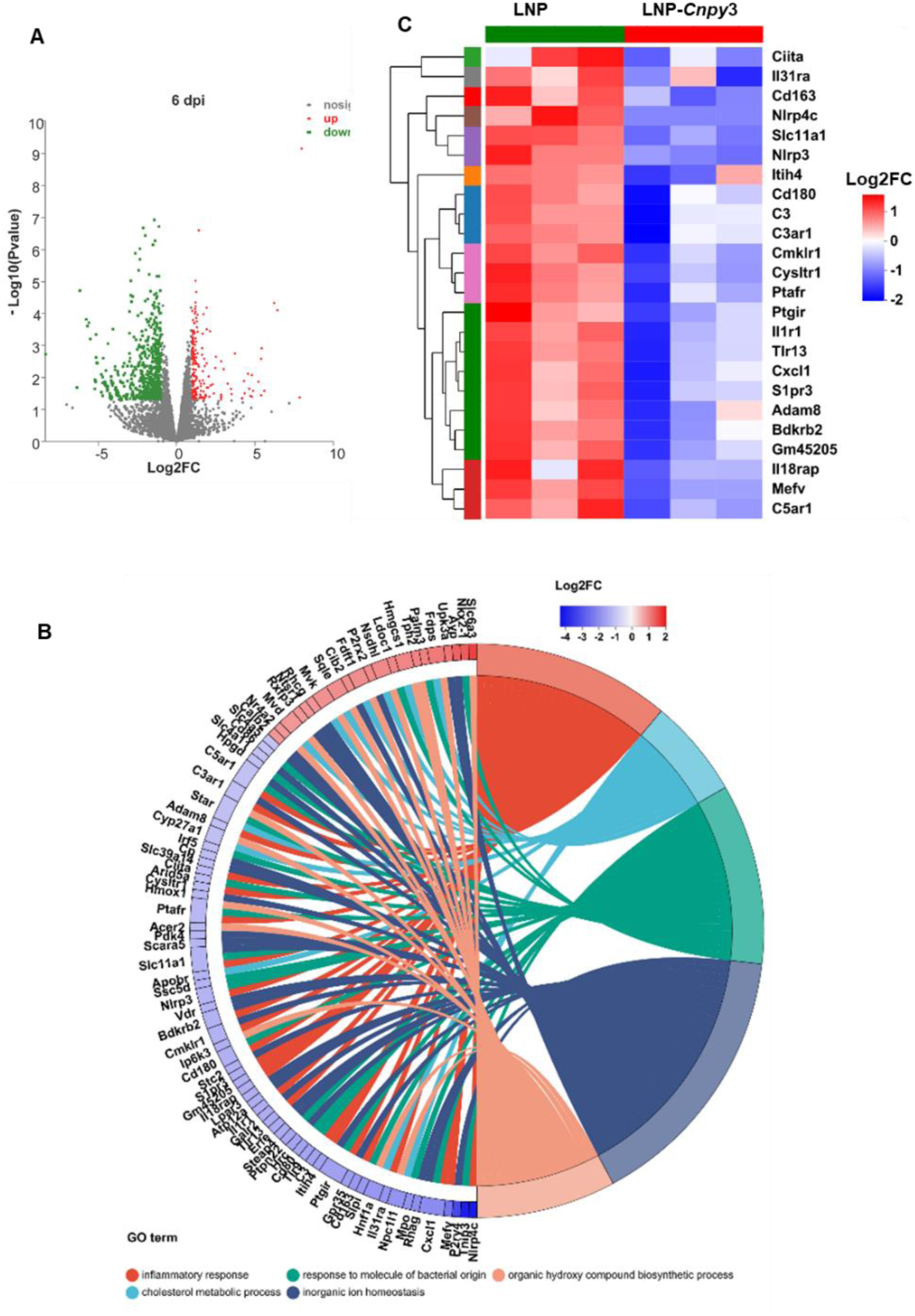
Transcriptomics analysis in later-stage flavivirus infection. (A) Volcano plot based on differentially expressed genes in samples. RNA-Seq analysis of brains from 3-day-old BALB/c mice brains infected with DENV-2 and treated with either LNP-*Cnpy3* or LNP alone on 6 days post-infection (dpi). Red and blue colors represent for up- and down-regulation of individual members, respectively. (B) Ontology (GO) enrichment analysis of DEGs. Chords represent a detailed relationship between the expression levels of DEGs (left semicircle perimeter) and their enriched GO pathways (right semicircle perimeter). For each gene, the expression value (log2 FC) of DEGs in brain tissue is shown by colored rectangles. (C) Genes of related to inflammatory response are presented by heatmap based on gene expression value. log2 FC are color coded as indicated.

### Supplementary Table

**Table S1.**
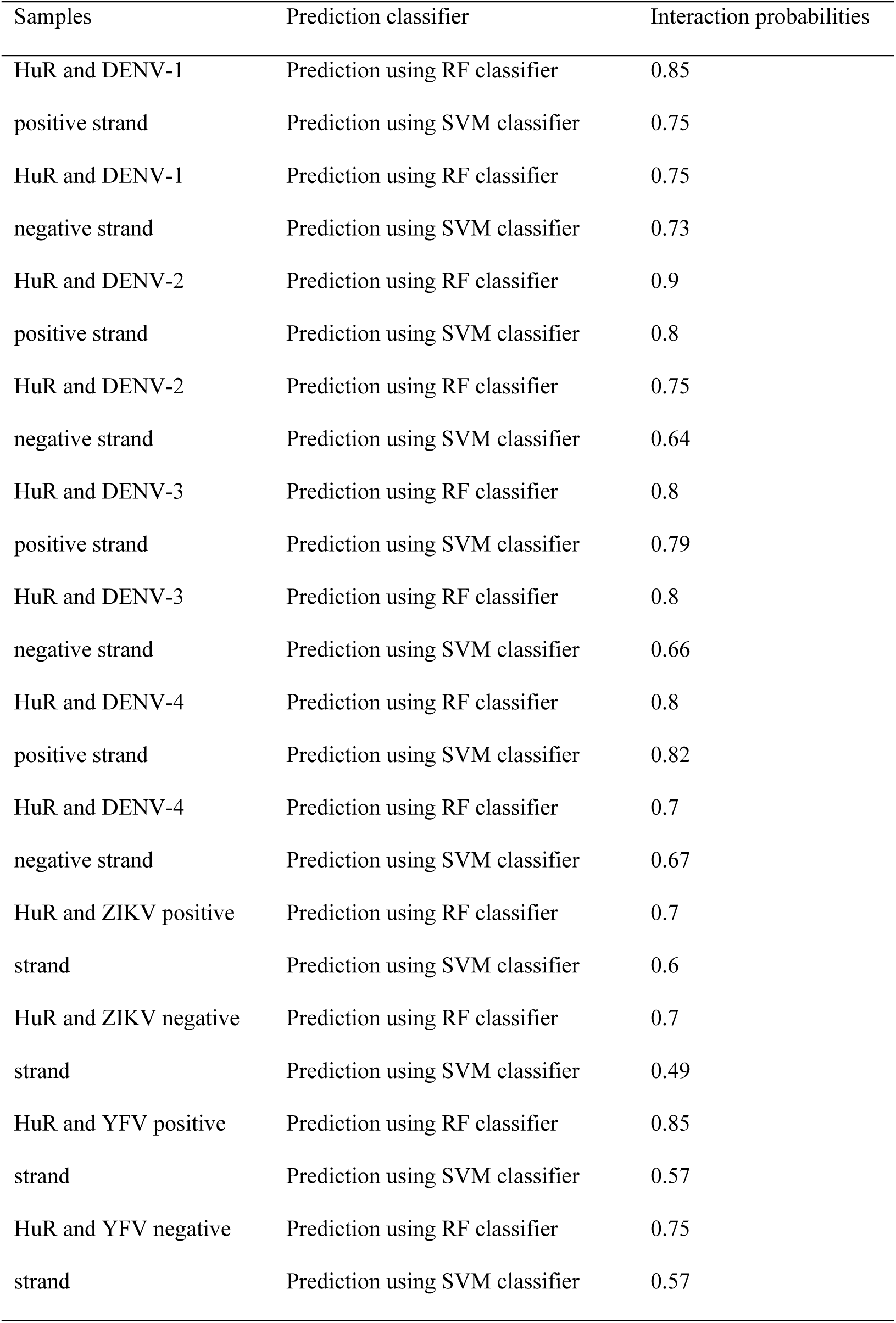
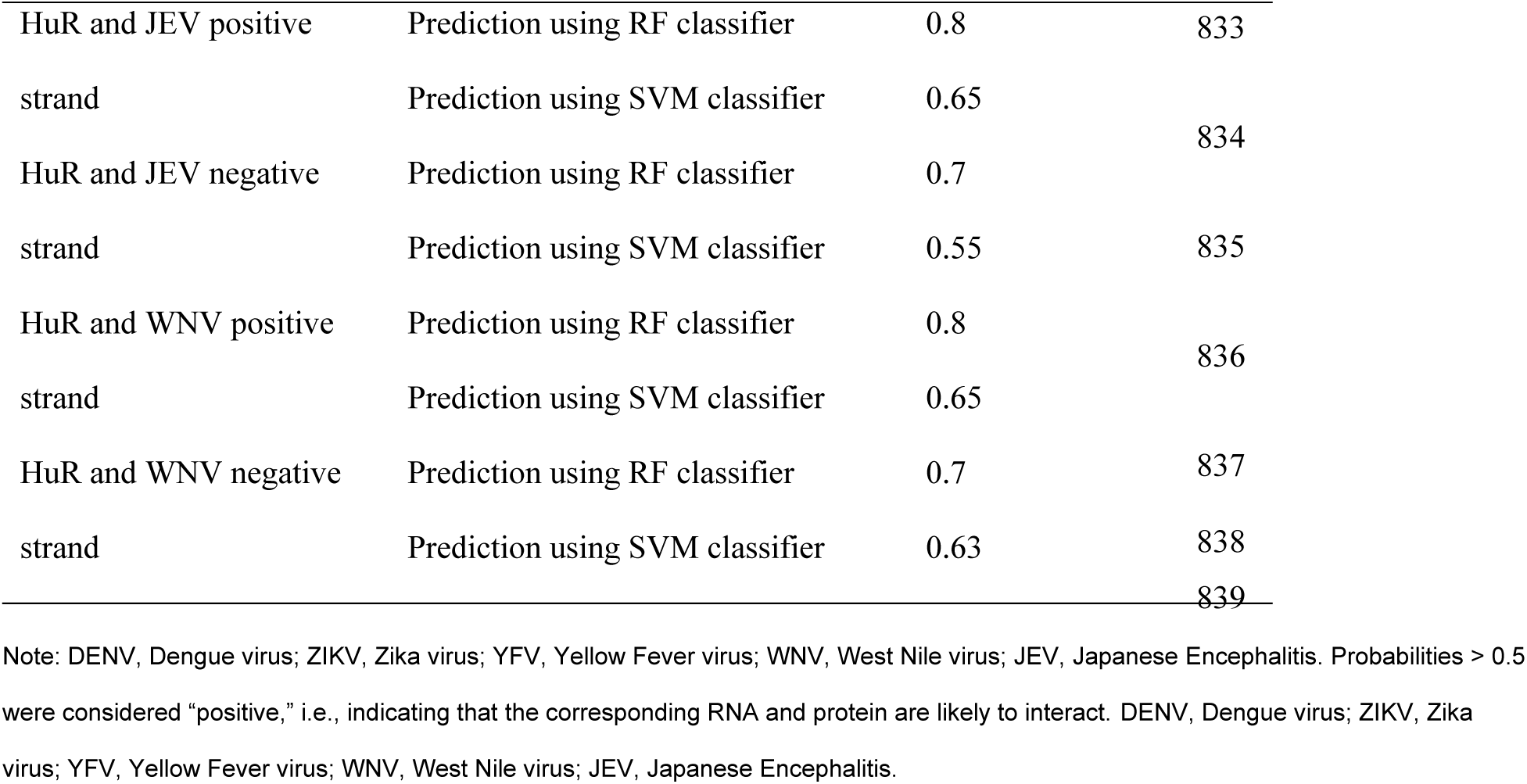
The interaction probabilities between HuR protein and Flavivirus.

### Supplementary Data

Data files S1 Primers, probes and RNA sequences were used for this paper.

